# Synaptic microRNA dysregulation orchestrates aberrant glutamatergic transmission in a human neuronal model of 22q11.2 deletion syndrome

**DOI:** 10.64898/2026.10.13.758925

**Authors:** Weibo Niu, Jidong Guo, Ximing Ran, David Parker, Xiangru Li, Shaojun Yu, Sidney Imes, Ryan Purcell, Christina Michalski, Eric Guo, Sophia Y. Jiang, Ethan Xu, Gary J Bassell, Jennifer G Mulle, Opal Y. Ousley, Bradley Pearce, Yue Feng, Carrie E. Bearden, Elaine F. Walker, Joseph F. Cubells, Zhaohui Qin, Erica Duncan, Zhexing Wen

## Abstract

22q11.2 Deletion Syndrome (22q11DS), the most prevalent chromosomal microdeletion disorder in humans (incidence ∼1 in 4,000 live births), arises from hemizygous deletions of a 1.5-3.0 Mb region on chromosome 22. Notably, 20-30% of individuals with 22q11DS develop schizophrenia (SCZ) by early adulthood, making 22q11DS one of the strongest genetic risk factors for SCZ. However, the mechanisms by which the 22q11.2 deletion perturbs neuronal function and drive psychotic manifestations in 22q11DS remain poorly defined. In this study, we generated human 2D cortical neurons and 3D cortical organoids from induced pluripotent stem cells (iPSCs) derived from clinically well-characterized individuals with 22q11DS. Cellular and electrophysiological analyses revealed that 22q11DS neurons exhibit hyperexcitability, reduced glutamatergic synaptic transmission, and impaired homeostatic synaptic plasticity, which are pathophysiological hallmarks that may be relevant to the clinical, cognitive, and psychophysiological profiles of patients. Transcriptomic profiling of neurons and synaptosomes from 22q11DS cortical organoids uncovered dysregulated immune pathways and synaptic gene networks. Integrative mRNA-microRNA network analysis identified synergistic downregulation of hsa-miR-185-5p and hsa-miR-128-3p at synapses, driving aberrant regulation of immune and synaptic transcripts. Experimental knockdown of these microRNAs in control neurons recapitulated the molecular and functional phenotypes of 22q11DS neurons, thereby establishing a mechanistic link to synaptic dysfunction and impaired plasticity. Collectively, these findings delineate a multimodal mechanistic framework that bridges genetic vulnerability, synaptic pathology, and psychosis penetrance in 22q11DS, positioning dysregulated synaptic microRNAs as nodal regulators of neurodevelopmental networks and potential therapeutic targets.

## INTRODUCTION

Schizophrenia (SCZ) is a severe and disabling neuropsychiatric disorder that affects approximately 1% of the population worldwide and is characterized by impairments in cognition, perception, and affective regulation ^1^. Despite extensive research, the biological mechanisms that give rise to SCZ remain incompletely understood, largely due to the disorder’s complex genetic architecture and polygenic inheritance. Genome-wide association studies and large-scale sequencing efforts have identified hundreds of common and rare variants that contribute to SCZ risk ^2,3^, yet the effect size of most variants is modest and the mechanisms by which genetic risk converges on neural dysfunction remain elusive. One promising strategy to advance mechanistic understanding is to focus on rare, highly penetrant genetic syndromes that confer disproportionately high risk for SCZ. These syndromes provide powerful entry points for linking defined genetic lesions to cellular, synaptic, and circuit-level abnormalities relevant to psychosis.

Among such syndromes, 22q11.2 Deletion Syndrome (22q11DS) stands out as one of the most compelling genetic models for SCZ. Historically, 22q11DS has also been referred to as DiGeorge syndrome or velocardiofacial syndrome. 22q11DS is the most common chromosomal microdeletion disorder in humans, arising from hemizygous deletion of a 1.5–3.0 Mb region on chromosome 22 and occurring in approximately 1 in 4,000 live births ^4^. The syndrome presents with a broad spectrum of clinical manifestations, including congenital anomalies, intellectual disability, and elevated risk for psychiatric disorders. Notably, 20–30% of individuals with 22q11DS develop SCZ by early adulthood SCZ ^5–8^, making it one of the strongest known genetic risk factors for psychosis. This remarkable penetrance positions 22q11DS as a uniquely tractable framework for investigating how defined genetic lesions translate into disease-relevant neuronal phenotypes and ultimately contribute to psychotic illness.

Although neuroimaging and clinical studies in 22q11DS cohorts have revealed alterations in cortical development, synaptic architecture, and cognitive trajectories ^9,10^, the pathophysiological mechanisms that bridge the 22q11.2 deletion to psychosis vulnerability remain poorly defined. A converging body of evidence suggests that synaptic dysfunction, particularly impaired glutamatergic transmission and maladaptive plasticity, is central to the pathogenesis of SCZ ^11,12^. Cortical excitatory neurons, which form the backbone of cognitive and perceptual processing, appear to be particularly vulnerable to genetic risk perturbations. Yet the molecular pathways that disrupt excitatory synaptic function in 22q11DS are incompletely understood.

Human induced pluripotent stem cells (iPSCs) have emerged as a robust and versatile platform for dissecting the molecular and cellular underpinnings of complex neuropsychiatric disorders, including SCZ^13–21^. In the present study, we leveraged iPSC-derived 2D cortical neurons and 3D cortical organoids from clinically well-characterized 22q11DS patients to interrogate the synaptic consequences of the 22q11.2 microdeletion and elucidate the underlying molecular mechanisms. Strikingly, electrophysiological analyses identified convergent neuronal phenotypes in 22q11DS, including hyperexcitability, reduced glutamatergic transmission, and impaired homeostatic plasticity. Further integrative transcriptomic analysis on synaptosomes revealed aberrant immune signaling and synaptic gene networks, largely driven by the synergistic downregulation of microRNAs hsa-miR-185-5p and hsa-miR-128-3p. Functional perturbation experiments demonstrated that knockdown of these miRNAs in healthy control neurons recapitulated 22q11DS-associated synaptic phenotypes, thereby establishing a causal mechanistic link between miRNA dysregulation, synaptic dysfunction, and vulnerability to psychosis. Together, these findings delineate a mechanistic framework that connects a defined genetic lesion to synaptic pathology through dysregulated miRNA networks, highlighting novel nodal regulators of neurodevelopmental transcriptomic programs. More broadly, this work underscores the power of human iPSC-derived neuronal models to illuminate the molecular substrates of psychiatric disease risk and to identify potential therapeutic targets for 22q11DS and schizophrenia.

## MATERIALS AND METHODS

### Psychosis and cognition assessment

For comprehensive details on the collection methods of human phenotyping data, including inclusion and exclusion criteria and the complete battery of measurements, please refer to our previous study^22^. Subjects completed assessments of prodromal symptoms severity from the Structured Interview for Psychosis-risk Syndromes (Positive, Negative, Disorganized and General)^23^, and Cognition (Average t-score from MATRICS subscales: Speed of Processing, Attention/Vigilance, Working Memory, Verbal Learning, Visual Learning, and Reasoning and Problem Solving)^24,25^.

### Auditory mismatch negativity response

Subjects were first screened for hearing acuity in both ears using a sound pressure audiometer (Model GS1710, Grason-Stadler, Eden Prairie, MN). For inclusion, subjects had to be able to detect tones bilaterally at a threshold of 40dB[A] at frequencies of 250, 500, 1000, 2000, and 4000 Hz. EEG-based evoked potentials were recorded using a Brain Vision EEG recording system (Brain Vision LLC, Morrisville, NC) from Fz and Cz electrodes with Fcz as the reference and Fpz as the ground electrode, with a sampling rate of 1000 Hz. Additional electrodes were placed both above and lateral to the right eye to detect extra-ocular movements. The Event-Related-Potential (ERP) paradigm was presented using Presentation software (Neurobehavioral Systems Inc., Albany, CA), as implemented in the North American Prodromal Longitudinal Studies (NAPLS) consortium study of individuals at high risk for psychosis^26,27^. The paradigm consisted of a pseudorandom sequence of frequent (85%) standard tones (Standards; 633 Hz, 50 ms duration) and infrequent (15%) deviant tones that differed in frequency pitch (Frequency Deviant; 1000 Hz, 50 ms), duration (Duration Deviant; 633 Hz, 100 ms) and both (Double Deviant; 1000 Hz, 100 ms)^28–30^. 1794 stimuli were presented at 85 dB with a stimulus onset asynchrony of 510 ms and a recording time of about 1 hour. During the MMN paradigm, participants were instructed to attend to an unrelated visual task. Raw EEG data were processed using Brain Vision Analyzer software (Brain Vision LLC, Morrisville, NC). EEG files were reviewed to flag artifacts. EEG files were processed through a scripted pipeline performing bandpass filtering (.01-30 Hz), eye movement/blink correction ocular correction method^31^, epoching, baseline correction, artifact rejection (+75 microvolts at sensors Fz and Cz), and averaging of trials for each stimulus type to generate ERPs^22^. Data from included trials were averaged for each participant to create a grand average ERP for each trial type for each sensor Fz and Cz. Each ERP was baseline adjusted using the 50-ms prestimulus period. Mismatch responses were generated by subtracted the ERP response to Standard tones from the ERP response to deviant tones. MMN neuronal correlational analyses were completed on the MMN response from sensor Cz to the double deviant during the 230-260 ms window. These three responses were selected for their representation of different analytical levels, aiming to shed light on the intricate relationship between basic neuronal function and human-level responses.

### Subjects and iPSC generation

Peripheral blood samples (10-15 mL) from four 22q11DS patients and four controls (Supplementary Table 1) were collected at the Atlanta Veterans Administration Health Care System, Atlanta, Georgia, USA, followed by isolation of peripheral blood mononuclear cells (PBMCs). Induced pluripotent stem cells (iPSCs) were further generated using the CytoTune™-iPS 2.0 Sendai Reprogramming Kit (ThermoFisher) in accordance with the manufacturer’s instructions. The iPSCs were fully characterized, including karyotyping, genotyping, assessment of pluripotency marker expression, and evaluation of in vitro differentiation capability. One of the control iPSC lines (C19) was used to generate an isogenic 22q11DS line (1F3) through the introduction of a 22q11.2 deletion via CRISPR-Cas9 gene editing, which was performed by the laboratory of Dr. Ryan Purcell. The CRISPR-engineered isogenic 22q11.2 deletion line was generated and validated as previously described ^32^. Briefly, the expected deletion was confirmed by TaqMan copy-number assays, and genome-wide optical mapping confirmed the 22q11.2 deletion as the only new structural variant detected relative to the parental line. The study was approved by the Institutional Review Board (IRB; CR007-IRB00111990). Written informed consent was obtained from all subjects or their legal representatives prior to study participation.

### Generation of NGN2-induced cortical neurons from human iPSCs

Human iPSCs were dissociated using Accutase (ThermoFisher) and replated as single cells on Matrigel-coated 12-well plates at a density of 380,000 cells per well in mTeSR1 medium (StemCell Technologies). On the following day, cells were infected with pTet-O-NGN2-puro lentivirus and Fudelta GW-rtTA lentivirus (Addgene). When approximately 75% confluence was reached, NGN2 expression was induced by the addition of doxycycline (2 μg/mL; Sigma) in KSR medium, which consisted of 415 mL KnockOut DMEM (ThermoFisher), 75 mL KnockOut Serum Replacement (ThermoFisher), 5 mL GlutaMAX supplement (ThermoFisher), 5 mL MEM non-essential amino acids (ThermoFisher), and 0.5 mL beta-mercaptoethanol (ThermoFisher). On day 2, the medium was replaced with a 1:1 mixture of KSR and N2B medium supplemented with puromycin (5 μg/mL; ThermoFisher) and doxycycline (2 μg/mL). The N2B medium was composed of DMEM/F12 (ThermoFisher), 1X GlutaMAX (ThermoFisher), 0.3% dextrose (Sigma), and 1X N2 supplement (ThermoFisher). On day 3, the medium was replaced with N2B medium supplemented with 1X B27 (ThermoFisher), puromycin (5 μg/mL), and doxycycline (2 μg/mL). On day 4, following puromycin selection, NGN2-induced neurons were dissociated using Accutase and cryopreserved. These day 4 neuronal stocks were subsequently thawed and replated on Matrigel-coated plates at a density of 2 million cells per plate in NBM medium for downstream experiments. The NBM medium was composed of 485 mL Neurobasal Medium (ThermoFisher), 5 mL GlutaMAX, 7.5 mL 20% dextrose, 2.5 mL MEM NEAA, 10 mL B27, 10 ng/mL recombinant human BDNF (PeproTech), 10 ng/mL recombinant human GDNF (PeproTech), 5 μg/mL puromycin, and 2 μg/mL doxycycline. Medium changes were performed twice per week. NGN2-induced neurons were maintained for 2-8 weeks after differentiation, depending on the downstream experiment. For electrophysiological experiments, NGN2-induced neurons were co-cultured with mouse astrocytes throughout neuronal maturation until electrophysiological recordings were performed. Human iPSCs were differentiated into NGN2-induced cortical-like neurons using an NGN2-based differentiation protocol and are hereafter referred to as cortical neurons for simplicity.

### Generation of cortical organoids from human iPSCs

All induced pluripotent stem cell (iPSC) lines were maintained on Matrigel-coated (Corning) plates and were cultured in mTeSR™1 medium (Stemcell Technologies). Cortical organoids were differentiated from iPSCs using a previously described protocol with modifications^33–37^. Briefly, intact human iPSC colonies were collected using Collagenase IV (Gibco) and transferred to 10 cm suspension culture dishes. Embryoid body (EB) media were used, with daily media changes. The EB medium was composed of DMEM/F12 (Corning), 20% KnockOut Serum Replacement (ThermoFisher), 1X GlutaMAX™ (ThermoFisher), 1X MEM non-essential amino acids solution (Invitrogen), 0.1 mM beta-mercaptoethanol (ThermoFisher), 2 μM A-83, and 2 μM Dorsomorphin dihydrochloride (Tocris/Fisher). On day 6, the EB media were replaced with neural medium (NM) composed of Neurobasal (Invitrogen), 1X B-27 serum substitute without vitamin A (Invitrogen), 1X GlutaMAX™, and 100 U/ml penicillin-streptomycin (Invitrogen). This NM was supplemented with 20 ng/ml bFGF (Peprotech) and 20 ng/ml EGF (Peprotech). Media were changed daily for the first 10 days, and every other day for the next 9 days. On day 25, to induce neuronal differentiation, bFGF and EGF were replaced with 20 ng/ml BDNF (Peprotech), and NM supplemented with BDNF was used with media changes every other day. On day 43, the media were switched to growth factor-free NM, which consisted of Neurobasal, 1X B-27 serum substitute without vitamin A, 1X GlutaMAX™, and 100 U/ml penicillin-streptomycin, and were changed every four days. From day 70 onward, cortical organoids were continuously maintained in NM containing 1X B-27 supplement with vitamin A (ThermoFisher), with medium changes every four days, until day 98.

### Immunocytochemistry

Cells cultured on coverslips were fixed in 4% paraformaldehyde (PFA; VWR) for 15 minutes at room temperature. Samples were then permeabilized and blocked with 0.5% Triton X-100 (Sigma) and 10% normal donkey serum (Sigma) in PBS for 20 minutes, as previously described^38,39^. Following this, samples were incubated with primary antibodies overnight at 4 °C, and subsequently with secondary antibodies for 2 hours at room temperature. The following primary antibodies were applied: anti-BRN2 (rabbit, 1:500; Proteintech, 14596-1-AP) and anti-MAP2 (chicken, 1:500; Novus, NB300-213). Secondary antibodies used for immunostaining included donkey anti-rabbit Alexa Fluor 568 (1:1000; ThermoFisher #A10042), anti-chicken Cy5 (1:1000; Jackson ImmunoResearch #703-175-155), and Hoechst 33342 (1:2000; Thermo Fisher, 62249). All antibodies were prepared in PBS containing 0.25% Triton X-100 and 10% donkey serum. After incubation with secondary antibodies, slides were washed three times with PBS, each wash lasting 10 minutes. A drop of Gold antifade reagent was applied to the coverslip, which was then mounted onto the slide. All images were captured using a Nikon Eclipse Ti-E microscope. Quantification of marker-positive cells was performed using ImageJ software (National Institutes of Health).

### Electrophysiology

Whole-cell patch clamp recordings were performed on neurons 2 to 8 weeks after differentiation. Coverslips containing neuronal cultures were transferred to a Warner recording chamber (RC-26GLP), which was continuously perfused with HEPES-based buffer at a rate of approximately 1 mL/min and maintained at 32 °C using a Warner TC-324C inline heater. The composition of the HEPES buffer was (in mM): NaCl 128, KCl 5, HEPES 25, CaCl₂ 2, MgCl₂ 1, and glucose 30; the pH was adjusted to 7.30 and the osmolarity was approximately 300 mOsm. Individual neurons were visualized on a Windows PC monitor using a Q-Electro CCD camera connected to a Leica DMLFS microscope equipped with differential interference contrast (DIC) optics and infrared (IR) illumination. Patch electrodes were pulled from borosilicate glass using a Sutter P-1000 Flaming/Brown micropipette puller, and had resistances ranging from 3 to 5 MΩ. The internal patch solution consisted of (in mM): 130 K-Gluconate, 8 KCl, 20 HEPES, 8 phosphocreatine, 1 EGTA, 4 Mg-ATP, and 0.3 Na-GTP. The solution was adjusted to pH 7.3 with KOH and had an osmolarity of 280–290 mOsm. For some recordings, 0.3% biocytin was included in the patch solution for post hoc morphological analysis. Recordings were conducted using a Multiclamp 700B amplifier (Molecular Devices) controlled by pClamp 11.0 software and an Axon Digidata 1550B A/D interface (Molecular Devices). Whole-cell access resistances ranged from 5 to 20 MΩ and were routinely monitored throughout each recording; recordings showing a change in access resistance greater than 15% were excluded from analysis. The membrane potential of neurons was held at –60 mV, and only neurons maintaining a stable resting membrane potential more negative than –40 mV were included for analysis. Intrinsic membrane properties were assessed in current-clamp mode as previously described^40,41^. Briefly, a series of hyperpolarizing and depolarizing current step injections (1 s duration) was applied to evaluate membrane currents and action potential firing patterns. A linear ramp protocol (250 ms duration) was also employed to elicit a single action potential for analysis of action potential kinetics.

### Recording of excitatory synaptic transmission and homeostatic plasticity

Spontaneous excitatory postsynaptic currents (sEPSCs) were recorded by holding the neuronal membrane potential at –60 mV. Spontaneous activity was continuously recorded for 2 minutes, and individual events were subsequently detected offline using MiniAnalysis software version 6.0 (Synaptosoft Inc., Decatur, GA). To induce AMPA receptor-dependent homeostatic plasticity, neuronal cultures were treated with tetrodotoxin (TTX; 1 µM) and DNQX (20 µM) for 24 hours. For recording, coverslips were removed from 24-well plates and transferred to the recording chamber. Cultures were perfused with HEPES-based buffer containing TTX (1 µM) and the GABA_A receptor antagonist SR95531 (5 µM) for at least 30 minutes to ensure complete washout of DNQX. The composition of the HEPES buffer was (in mM): NaCl 140, KCl 5, HEPES 10, CaCl₂ 3, MgCl₂ 1, and glucose 10; the pH was adjusted to 7.30, and the osmolarity was ∼310 mOsm. The internal solution used for patch clamp recordings contained (in mM): CsMeSO₄ 138, MgCl₂ 2, HEPES 10, EGTA 0.25, sodium phosphocreatine 5, MgATP 4, NaGTP 0.3, NaCl 8, and QX-314 2. The pH was adjusted to 7.3 with CsOH, and the osmolarity was ∼290 mOsm. For each neuron, a continuous 5-minute recording was obtained. All miniature EPSC events were analyzed using MiniAnalysis version 6.0.

### Knockdown of miRNAs in a control iPSC line

The hsa-miR-185-5p and hsa-miR-128-3p knockdown lentiviral vectors and their corresponding lentiviral particles were designed and produced by VectorBuilder. The shRNA targeting hsa-miR-185-5p spans residues 2566-2615 and is 50 nucleotides in length. The sequence is TCAGGAACTGCCTTTCTCTCCACTCGAGTGGAGAGAAAGGCAGTTCCTGA. The shRNA targeting hsa-miR-128-3p spans residues 2566-2613 and is 48 nucleotides in length. The sequence is: AAAGAGACCGGTTCACTGTGACTCGAGTCACAGTGAACCGGTCTCTTT. Lentivirus was added to Day 28 cortical neurons (004 cell line) at a multiplicity of infection (MOI) of 5. Neuron samples were collected on Day 42. Neurons were homogenized in TRIzol Reagent (Invitrogen, 15596018), and RNA was isolated according to the manufacturer’s instructions using the Direct-zol™ RNA Miniprep Kit (Zymo, R2052). miRNA cDNA was synthesized using the TaqMan™ Advanced miRNA cDNA Synthesis Kit (Cat. No. A28007) according to the manufacturer’s instructions. The expression of hsa-miR-185-5p and hsa-miR-128-3p was then assessed by qPCR using the TaqMan™ Fast Advanced Master Mix (Cat. No. 4444557). Each reaction was performed in triplicate and analyzed using the ΔΔCt method, with hsa-miR-186-5p^42^ (ThermoFisher; 477940_mir) serving as the normalization control. Expression of hsa-miR-185-5p (ThermoFisher; 477939_mir) and hsa-miR-128-3p (ThermoFisher; 477892_mir) was measured using the TaqMan™ Advanced miRNA Assays.

### RNA isolation, RNA sequencing, and miRNA sequencing

Synaptosomes from cortical organoids were extracted using the Syn-PER Synaptic Protein Extraction Reagent (Thermo, 87793). Briefly, cortical organoids were homogenized in ice-cold Syn-PER reagent and centrifuged at 1,200 × g for 10 min at 4°C to remove cellular debris. The resulting supernatant was then centrifuged at 15,000 × g for 20 min at 4°C, and the synaptosome pellet was collected for subsequent RNA isolation. Neurons or synaptosomes were homogenized in TRIzol Reagent (Invitrogen, 15596018), and RNA was isolated according to the manufacturer’s instructions using the Direct-zol™ RNA Miniprep Kit (Zymo, R2052).

Total RNA was quantified using the Qubit 2.0 Fluorometer (ThermoFisher Scientific, Waltham, MA, USA) and assessed for integrity with the 4200 TapeStation (Agilent Technologies, Palo Alto, CA, USA). Strand-specific libraries were prepared using the NEBNext Ultra II Directional RNA Library Prep Kit for Illumina (NEB, Ipswich, MA, USA), following the manufacturer’s instructions. RNA was fragmented at 94 °C for 8 minutes, and first- and second-strand cDNA synthesis was performed, with dUTP incorporated during second-strand synthesis to maintain strand specificity. After 3′ end adenylation, adapter ligation, and limited-cycle PCR amplification, libraries were validated using the Agilent TapeStation and quantified by Qubit 2.0 (ThermoFisher Scientific) and qPCR (KAPA Biosystems, Wilmington, MA, USA). Libraries were multiplexed, clustered onto a flowcell, and sequenced on the Illumina NovaSeq 6000 system using a 2 × 150 bp paired-end configuration, according to the manufacturer’s protocol. Image analysis and base calling were performed using the NovaSeq Control Software (Illumina, San Diego, CA, USA), and raw BCL files were converted to FASTQ format and demultiplexed with bcl2fastq v2.20 (Illumina), allowing one mismatch for index recognition.

Small RNA libraries were prepared using the NEBNext Small RNA Library Prep Kit (New England Biolabs, Ipswich, MA, USA) following the manufacturer’s protocol. Adapters were ligated to RNA with appropriate end structures, followed by reverse transcription and PCR amplification with indexed primers. Amplified cDNA was size-selected (∼145–160 bp) by PAGE, eluted, and concentrated by ethanol precipitation. Libraries were validated using the Agilent TapeStation (Agilent Technologies, Palo Alto, CA, USA) and quantified by Qubit 2.0 Fluorometer (ThermoFisher Scientific, Waltham, MA, USA) and qPCR (KAPA Biosystems, Wilmington, MA, USA). Sequencing was performed on the Illumina NovaSeq 6000 using a 2 × 150 bp paired-end configuration. Image analysis and base calling were conducted with NovaSeq Control Software, and FASTQ files were generated and demultiplexed using bcl2fastq v2.20 (Illumina, San Diego, CA, USA), allowing one mismatch in index recognition.

Libraries were aligned to the GRCh38 reference genome (Ensembl release-106 annotation) using HISAT2 v2.2^43^, which employs a hierarchical graph FM index for spliced alignment. The alignments (SAM) were sorted directly into BAM format with Samtools v1.20^44^. Gene-level quantification was performed by featureCounts v2.0.2^45^ in paired-end, stranded mode assigning reads to exonic features based on the Ensembl GT. Post-processing involved extracting gene IDs and raw counts, sorting by gene identifier, and joining to a pre-compiled Ensembl gene ID–to– gene name mapping (release 106). Differential expression analysis was performed using the DESeq2 package v1.26.0^46^ in R. Multiple testing correction was performed using the Benjamini-Hochberg procedure to control the false discovery rate (FDR). Genes with an FDR-adjusted *P* value < 0.05 and an absolute log2 fold change > 1.0 were defined as differentially expressed genes (DEGs) or differentially expressed miRNAs (DEMs). Gene set enrichment analysis was conducted using the R package gprofiler2 v0.2.2^47^, with multiple testing correction performed using the Benjamini–Hochberg procedure, and GO terms with an FDR-adjusted *P* value < 0.05 were considered significant. Protein-protein interaction networks were generated using the STRINGdb package^48^ with a confidence score threshold > 400. miRNA targets were predicted using the ENCORI database^49^, which integrates seven major algorithms: PITA, RNA22, miRmap, microT, miRanda, PicTar, and TargetScan. Predicted upregulated miRNAs were integrated with experimentally observed upregulated miRNAs to identify key regulators. A miRNA-gene regulatory network was then constructed at the pathway level. Pathway-level variation in gene expression was assessed via Gene Set Variation Analysis (GSVA) using the GSVA R package^50^, and GSVA scores were compared across sample groups using student t-test. All downstream statistical analyses and visualizations were performed using R (v4.2.0).

## RESULTS

### 22q11.2 deletion enhances neuronal excitability

To determine whether and how 22q11.2 deletion alters the synaptic function of cortical neurons which are fundamental to cognitive and perceptual processing and a key driver of SCZ pathogenesis^11,12^, we generated iPSC lines from four individuals with 22q11DS and four age- and sex-matched healthy controls after comprehensive clinical characterization (Fig. 1A-C; Supplementary Table 1). The psychosis-related symptoms and cognitive performance across multiple domains were assessed using the Structured Interview for Psychosis-risk Syndromes ^23^. Participants with 22q11DS exhibited significantly higher prodromal positive symptom severity scores (Fig. 1D), negative symptom severity scores (Fig. 1E), and disorganized symptoms (Fig. 1F) compared to controls. General symptom severity scores showed a trend towards an increase in the 22q11DS group (*P* = 0.0845; Fig. 1G). In addition to symptom severity, global functioning was significantly impaired, as indicated by a lower global assessment of functioning (GAF) score in the 22q11DS subjects compared to controls (Fig. 1H). Across multiple cognitive assessment domains individuals with 22q11DS were all significantly impaired in compared to controls and showed reduced attention (Fig. 1I), visual learning (Fig. 1J), speed of processing (Fig. 1K), working memory (Fig. 1L), as well as problem-solving ability (Fig. 1M).

**Figure 1.**
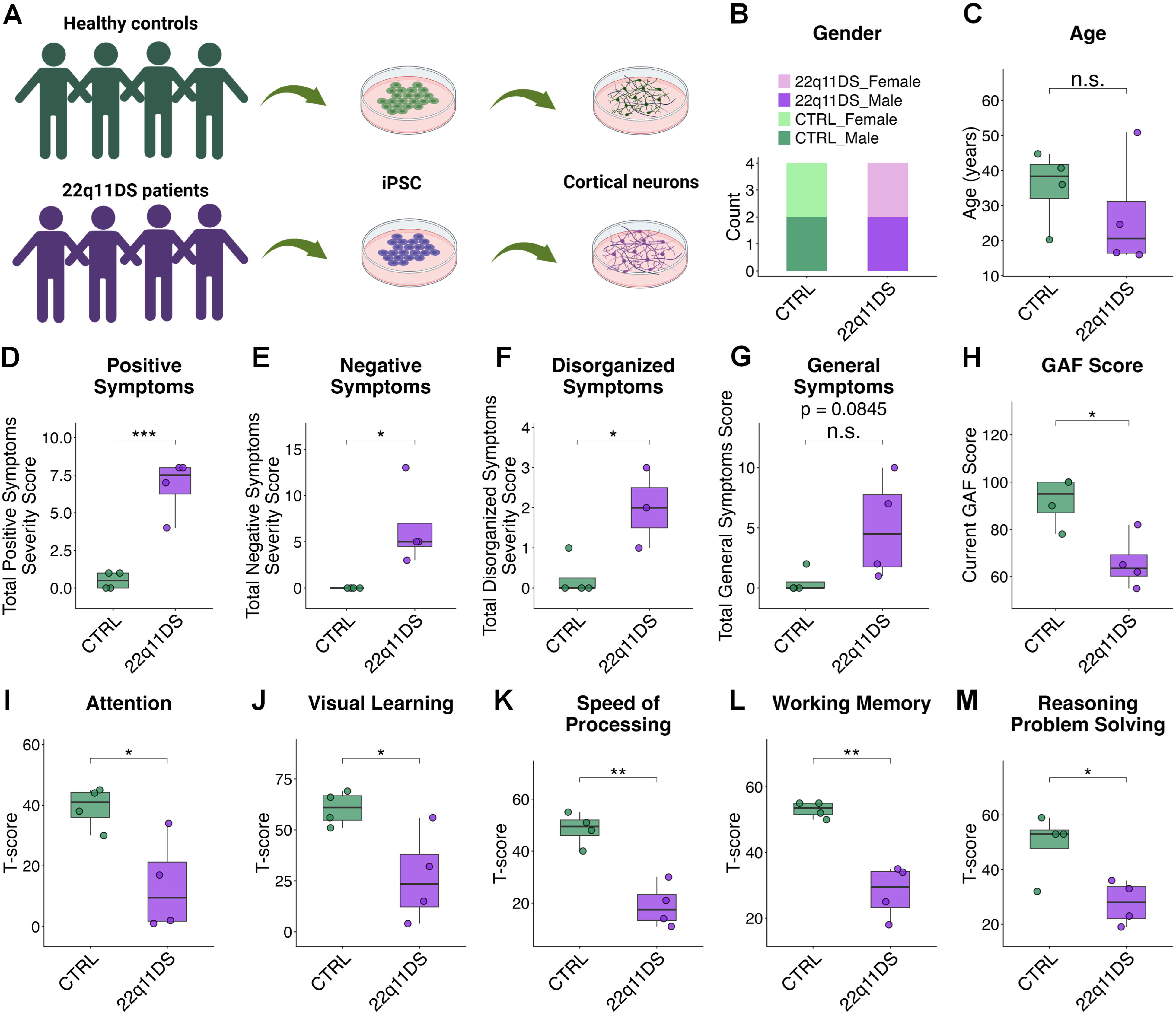
Clinical phenotypes related to psychosis and cognition in 22q11DS. (**A**) Schematic of experimental design. Induced pluripotent stem cells (iPSCs) derived from healthy controls (004, 006, 007, and 008) and 22q11DS patients (22q001, 22q002, 22q003, and 22q005) were differentiated into cortical neurons. (**B**) Sex distribution of control (CTRL) and 22q11.2 deletion syndrome (22q11DS) subjects. (**C**) Age information of CTRL and 22q11DS subjects. (**D-G**) Individuals with 22q11DS exhibited significantly higher positive (D), negative (E), and disorganized (F) symptom severity scores, while general symptoms (G) showed a trend toward significance (*P* = 0.0845). (**H**) Global assessment of functioning (GAF) score was significantly lower in the 22q11DS group. (**I-M**) Cognitive performance was impaired in 22q11DS individuals across several domains, including attention (I), visual learning (J), speed of processing (K), working memory (L), and reasoning/problem solving (M). Data are presented as individual data points with group means ± box plots (\**P* < 0.05, \*\**P* < 0.01, \*\*\**P* < 0.001; student’s t-test). n = 4 per group, except for disorganized symptom severity scores, where n = 3 for the 22q11DS group.

These iPSCs were subsequently directed to cortical excitatory neuron fate using an induced differentiation approach based on neurogenin-2 (NGN2) overexpression ^51^ (Fig. 1A). Immunostaining demonstrated BRN2 expression, validating the cortical neuronal identity of these differentiated cells (Supplementary Fig. 1A-B). We next quantified neurite morphology by labeling individual neurons with biocytin and performing Sholl analysis to assess process complexity and branching ^52^ (Fig. 2A). Neurons derived from 22q11DS exhibited a significantly increased number of distal branches compared to controls (Fig. 2B), indicating that the 22q11.2 deletion promotes enhanced neuritic arborization. We further assessed the membrane electrophysiological properties of 22q11DS neurons using whole-cell patch-clamp recordings performed on 2–8-week-old control and 22q11DS neurons (Fig. 2C-D, Supplementary Fig. 1C). Passive membrane properties, including resting membrane potential (Fig. 2E), input resistance (Fig. 2F), and membrane time constant (Tau) (Fig. 2G), did not differ significantly between the two groups. We next compared action potential (AP) kinetics and observed that 22q11DS neurons exhibited significantly lower AP thresholds and increased fast afterhyperpolarization amplitudes (fAHP) at 6 and 8 weeks post-differentiation relative to controls (Fig. 2H-I), indicating stage-specific hyperexcitability. Other AP parameters, including amplitude (Fig. 2J), rise time (Fig. 2K), decay time (Fig. 2L), and half-width (Fig. 2M), followed similar developmental trajectories in both groups.

**Figure 2.**
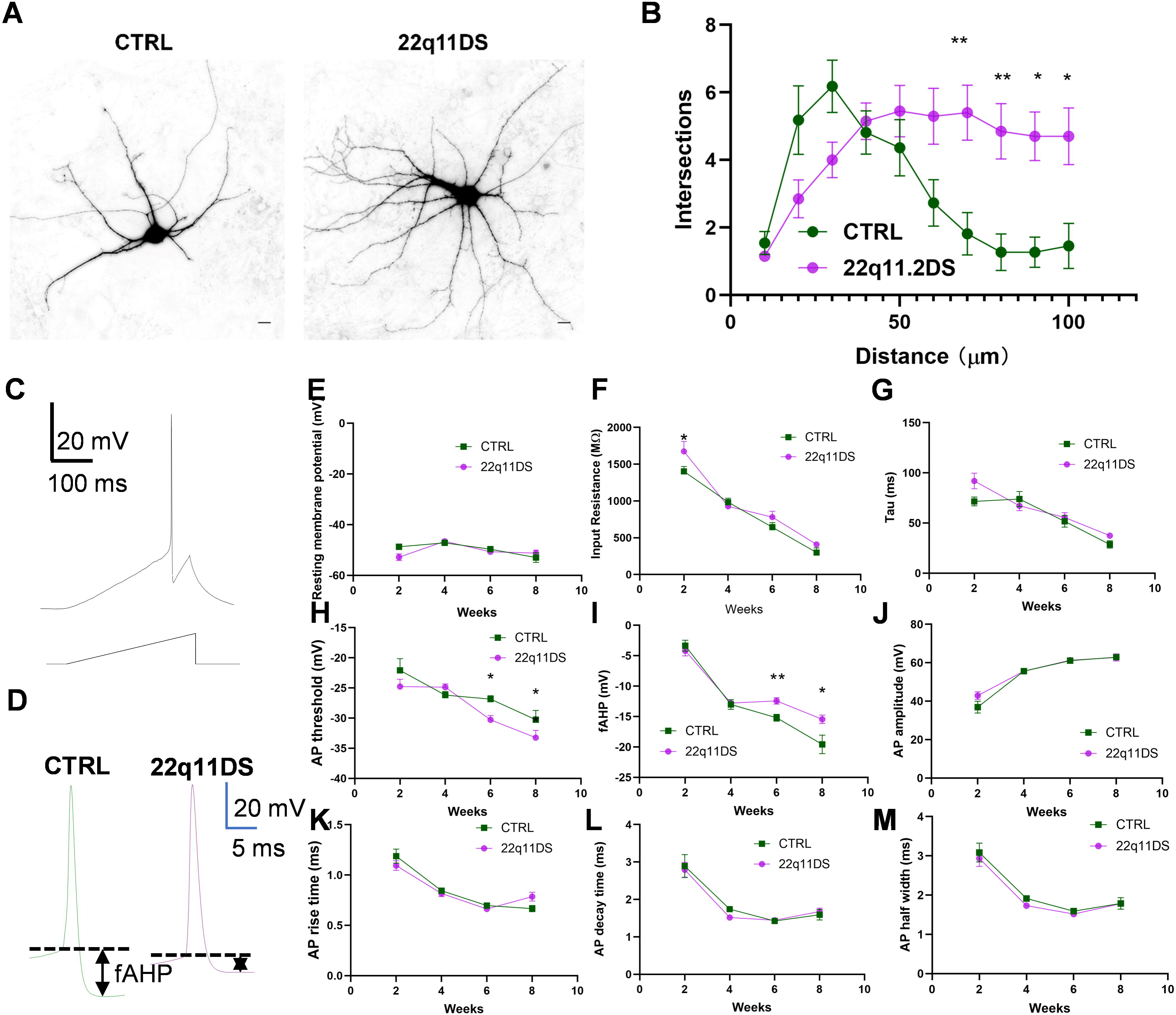
Morphological and intrinsic membrane properties of 22q11DS neurons. (**A-B**) Sample images (A) and Sholl analysis (B) of CTRL and 22q11DS neurons. Data are presented as mean ± s.e.m. (n = 11 CTRL neurons, n = 20 22q11DS neurons). \**P* < 0.05, \*\**P* < 0.01 (student’s t-test). (**C-M**) Electrophysiological properties of CTRL and 22q11DS cortical neurons were evaluated at 2-8 weeks after differentiation using patch-clamp recordings. Shown are sample tracings of action potentials in response to current steps (C) and first action potential (D), as well as quantifications of resting membrane potential (E), input resistance (F), membrane time constant (Tau) (G), AP threshold (H), fast afterhyperpolarization (fAHP; I), AP amplitude (J), AP rise time (K), AP decay time (L), and AP half-width (M). Data are presented as mean ± s.e.m. (2-week: n = 37 CTRL, n = 42 22q11DS; 4-week: n = 56 CTRL, n = 84 22q11DS; 6-week: n = 56 CTRL, n = 49 22q11DS; 8-week: n = 11 CTRL, n = 25 22q11DS; \**P* < 0.05, \*\**P* < 0.01, student’s t-test). AP: action potential.

### 22q11.2 deletion reduces excitatory synaptic transmission and disrupts homeostatic plasticity

To examine functional impacts of 22q11.2 deletion on glutamatergic synaptic transmission, we recorded and measured spontaneous excitatory postsynaptic currents (sEPSCs) from 6-week-old control and 22q11DS neurons (Fig. 3A). We found that the frequency of sEPSCs, but not the amplitude, was significantly lower for 22q11DS neurons compared to the controls (Fig. 3B-C), suggesting a decrease of excitatory synaptic transmission in 22q11DS neurons.

**Figure 3.**
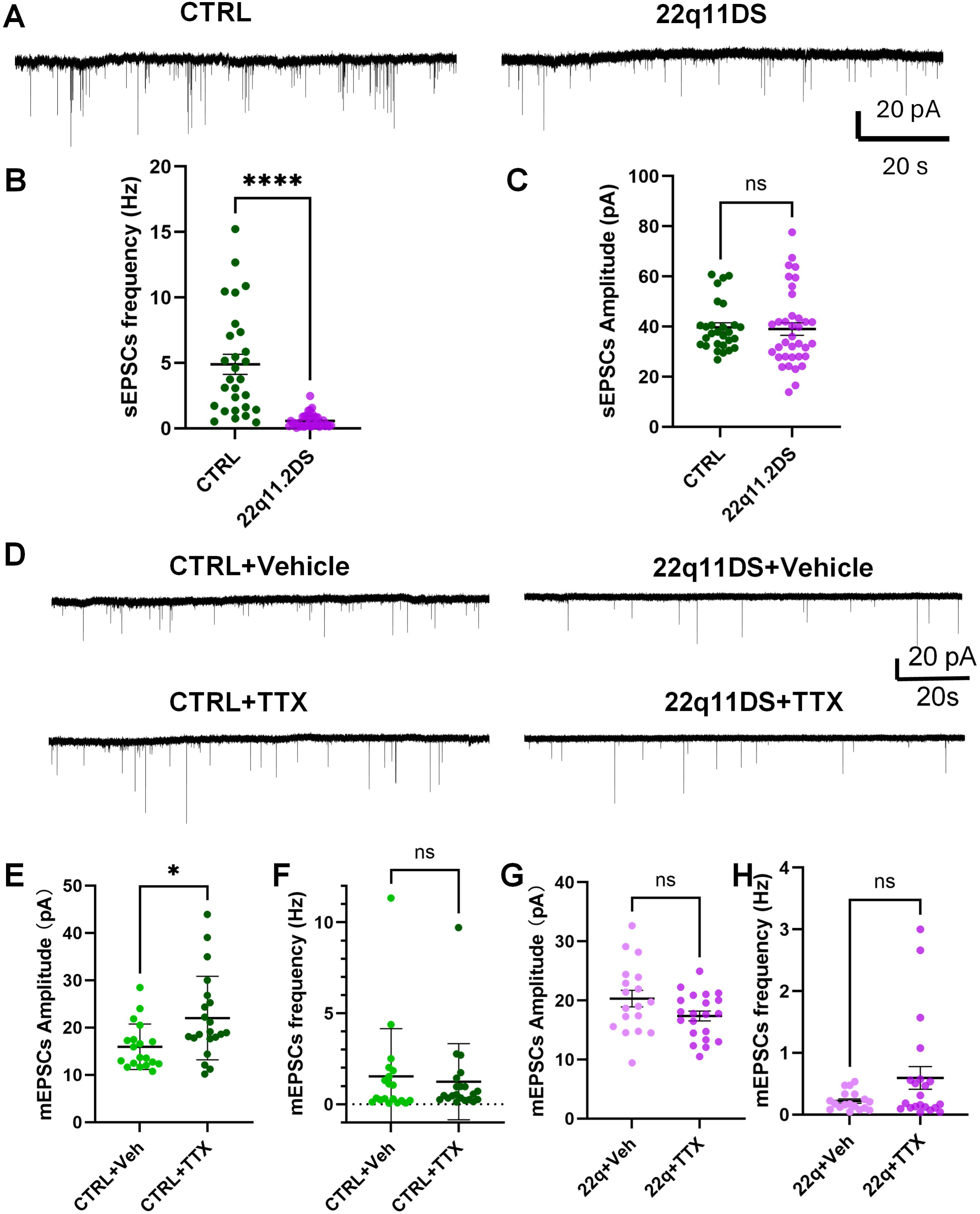
Reduced glutamate transmission and disrupted homeostatic plasticity in 22q11DS neurons. **(A-C)** Reduced glutamate transmission in 22q11DS neurons. Shown are sample traces (A) of sEPSCs recorded from CTRL and 22q11DS cortical neurons and quantification of the frequency (B) and amplitude (C) of sEPSCs (n = 28 CTRL, 36 22q11DS; \*\*\*\**P* < 0.0001, Student’s t-test). (**D-H**) Disruption of homeostatic plasticity in 22q11DS cortical neurons. Shown are representative traces (D) and quantification of mEPSCs amplitude (E and G) and frequency (F and H) recorded from CTRL and 22q11DS neurons (n = 19 CTRL + Veh, 21 CTRL + TTX, 18 22q11DS + Veh, 21 22q11DS + TTX; \**P* < 0.05, Student’s t-test).

Synaptic plasticity is critical for learning, memory, and cognition and is vulnerable to disruption in neurological and psychiatric disorders^53^. Homeostatic synaptic plasticity maintains neural circuit stability by adjusting synaptic strength in response to prolonged changes in neuronal activity^54,55^, and its disruption has been linked to psychotic disorders, such as SCZ^56,57^. To examine whether homeostatic synaptic plasticity is disrupted in 22q11DS cortical neurons, we inhibited neural network activity for 24 hours using tetrodotoxin (TTX; 1 µM) and the AMPA receptor antagonist DNQX (20 µM) in 6-week-old neurons. Miniature EPSCs (mEPSCs) were then recorded in the presence of TTX (1 µM) and the gamma-aminobutyric acid type A (GABA-A) receptor antagonist GABAzine (5 µM). Strikingly, in control neurons, synaptic silencing significantly increased mEPSC amplitude but did not affect mEPSC frequency (Fig. 3E-F), suggesting the induction of homeostatic plasticity. However, in 22q11DS neurons, TTX treatment resulted in a non-significant trend toward decreased mEPSC amplitude and increased mEPSC frequency (Fig. 3G-H), suggesting a disruption of homeostatic plasticity in 22q11DS neurons.

### 22q11.2 deletion perturbs gene modules associated with synaptic signaling and neuroimmune regulation

To gain molecular insight into the developmental and synaptic deficits associated with the 22q11.2 deletion, we performed bulk RNA-seq on control and 22q11DS cortical neurons at 6 weeks post-differentiation (Fig. 4A-C). We identified differentially expressed genes (DEGs) in 22q11DS neurons, including 891 upregulated genes and 87 downregulated genes (adjusted *P*-value < 0.05, absolute log2 fold-change > 1.0; Fig. 4A; Supplementary Table 2). To elucidate the molecular pathways affected by the 22q11.2 deletion, gene ontology (GO) analyses of DEGs were conducted (Fig. 4B), and revealed that upregulated DEGs in 22q11DS neurons are enriched in biological pathways associated with immune response, immune system process, inflammation response, cytokine production, and nervous system process (Fig. 4B left), while downregulated DEGs are involved in 22q11.2 copy number variation (CNV) syndrome, synaptic assembly, synaptic transmission (Fig. 4B right), which is consistent with the synaptic phenotypes identified in 22q11DS neurons (Fig. 3A-B). Gene interaction analysis revealed a highly interactive functional network composed of DEGs within the immune system process pathway (Fig. 4C).

**Figure 4.**
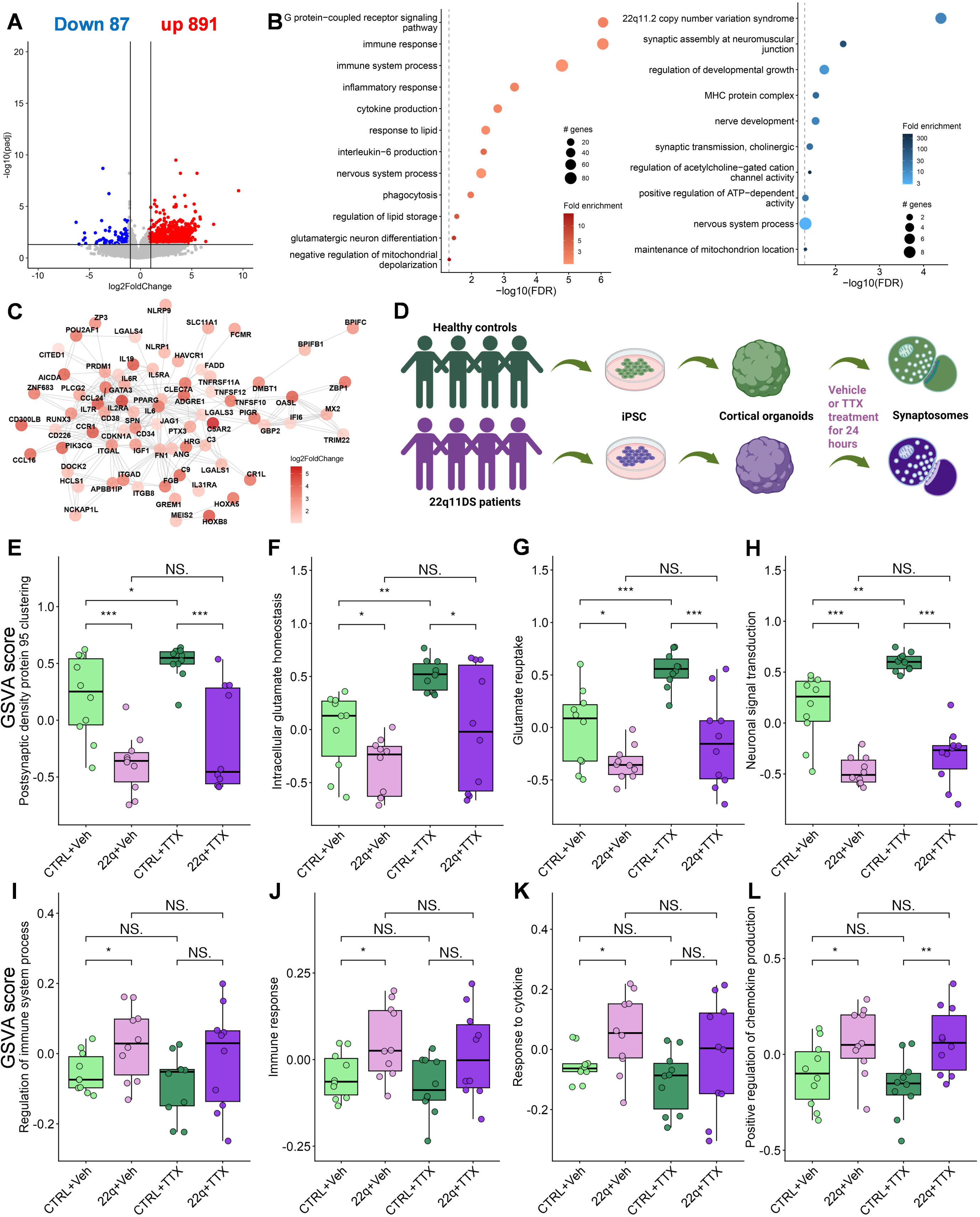
Transcriptomic analyses of 22q11DS neurons and synaptosomes. (**A**) Volcano plot shows differentially expressed genes (DEGs) in 22q11DS neurons derived from healthy controls (004, 006, 007, and 008) and 22q11DS patients (22q001, 22q002, 22q003, and 22q005), with two replicates per sample. Red and blue dots indicate 891 upregulated genes and 87 downregulated genes, respectively (adjusted *P*-value < 0.05, absolute log2 fold-change > 1.0). Grey dots represent genes with no significant differential expression. (**B**) Gene Ontology (GO) enrichment analysis of upregulated (right) and downregulated (left) DEGs in 22q11DS neurons compared with control neurons. Dot size represents the number of DEGs associated with each GO term, and color indicates fold enrichment. The x-axis represents −log10(FDR) (FDR < 0.05). (**C**) Gene interaction analysis of the immune system process pathway reveals a highly interactive functional network of DEGs in 22q11DS neurons. (**D**) Schematic of experimental design. Synaptosomes were isolated from day 98 cortical organoids derived from control iPSC lines (004, 006, 007, 008, and C19), 22q11DS patient iPSC lines (22q001, 22q002, 22q003, and 22q005), and the CRISPR-engineered C19 isogenic 22q11.2 deletion line (1F3) following 24-hour treatment with vehicle or TTX, with two replicates per sample. (**E-L**) Gene Set Variation Analysis (GSVA) of the transcriptome was performed on synaptosomes, including synapse-related gene sets postsynaptic density protein 95 clustering (E), intracellular glutamate homeostasis (F), glutamate reuptake (G), and neuronal signal transduction (H), as well as inflammation-related gene sets regulation of immune system process (I), immune response (J), response to cytokine (K), and positive regulation of chemokine production (L). Data are shown as box plots with individual data points. \**P* < 0.05, \*\**P* < 0.01, Student’s t-test.

Synapses are the fundamental units of neuronal communication and plasticity, and many disease-relevant molecular processes, including activity-dependent transcriptional remodeling, local translation, and receptor trafficking, are executed within synaptic compartments. Thus, bulk transcriptomic analyses of whole neurons may obscure subtle but functionally critical alterations that occur specifically at synapses. To address this limitation, we performed RNA-seq on synaptosomes isolated from cortical organoids. We generated 3D cortical organoids using an established protocol ^33–37^ from the four 22q11DS iPSC lines and four healthy iPSC lines, along with one pair of isogenic lines. The heterozygous isogenic line (1F3) was generated by introducing the 22q11.2 deletion into a wild-type control iPSC line (C19) using CRISPR/Cas9 genome-editing technology (Supplementary Table 1). We further inhibited neural network activity for 24 hours using tetrodotoxin (TTX; 1 µM) and the AMPA receptor antagonist DNQX (20 µM) in day 98 control and 22q11DS cortical organoids, followed by synaptosome isolation and RNA-seq (Fig. 4D). The low expression of 22q11.2 deletion genes in synaptosomes from 22q11DS iPSC-derived cortical organoids further confirm their genotypes (Supplementary Fig. 2). We identified numerous significant DEGs in 22q11DS synaptosomes, including 934 upregulated and 512 downregulated genes under basal condition (Supplementary Table 3), and 770 upregulated and 283 downregulated genes under homeostatic plasticity condition (Supplementary Table 3). We then conducted transcriptomic analyses and Gene Set Variation Analysis (GSVA)^50^ on synaptosomes across different conditions. Our results revealed that the GSVA scores for postsynaptic density protein 95 (PSD-95) clustering (Fig. 4E), intracellular glutamate homeostasis (Fig. 4F), glutamate reuptake (Fig. 4G), and neuronal signal transduction (Fig. 4H) were decreased in synaptosomes from 22q11DS organoids. Interestingly, these scores were significantly increased in control synaptosomes following TTX treatment, whereas no changes were observed in 22q11DS synaptosomes (Fig. 4E-H). These results align with our whole-cell patch-clamp recordings (Fig. 3), which demonstrated that homeostatic plasticity was induced in control but not in 22q11DS neurons, reinforcing the conclusion that the 22q11.2 deletion impairs excitatory synaptic transmission and disrupts homeostatic plasticity. Intriguingly, GSVA scores for regulation of the immune system process (Fig. 4I), immune response (Fig. 4J), response to cytokines (Fig. 4K), and positive regulation of chemokine production (Fig. 4L) were significantly upregulated in 22q11DS synaptosomes compared to controls, which is consistent with our bulk RNA-seq analysis of cortical neurons (Fig. 4B). These results indicate that the 22q11.2 deletion enhances synaptic neuroinflammatory signaling, a mechanism that may contribute to excitatory synaptic dysfunction in 22q11DS.

### Downregulated microRNAs reshape the synaptic transcriptome of 22q11DS

Accumulating data has implicated microRNAs (miRNAs) as critical mediators of neuronal development and synaptic function. These small non-coding RNAs regulate gene expression post-transcriptionally and shape broad transcriptomic networks underlying neuronal identity, synaptic signaling, and plasticity^58^. The 22q11.2 locus encodes several miRNAs, including hsa-miR-185-5p. Furthermore, *DGCR8* (DiGeorge syndrome critical region 8) and *RANBP1* (Ran-binding protein 1) are both located in the 22q11.2 deletion region and involved in miRNA biogenesis^59–62^. DGCR8 is an essential component of the microprocessor complex, working alongside Drosha to process primary miRNAs (pri-miRNAs) into precursor miRNAs (pre-miRNAs). This step is crucial in miRNA biogenesis, as it determines the maturation and functional availability of miRNAs for gene regulation^59,61^. RANBP1 is a key regulator of nuclear transport, interacting with Ran GTPase to facilitate nucleocytoplasmic trafficking. While not directly involved in miRNA processing, RANBP1 may influence miRNA biogenesis by regulating the nuclear export of pre-miRNAs, which are transported by Exportin-5 in a RanGTP-dependent manner^62,63^.

To investigate the mechanistic contribution of miRNAs to the transcriptional remodeling in 22q11DS synapses, we performed mRNA-seq and miRNA-seq on synaptosomes from control and 22q11DS organoids, followed by integrative transcriptome–miRNome analysis (Fig. 5A). We identified significantly differentially expressed miRNAs (DEMs), including 104 upregulated and 37 downregulated miRNAs (Supplementary Table 4). We utilized the predictive algorithms available in ENCORI^49^, including PITA, RNA22, miRmap, microT, miRanda, PicTar, and TargetScan, to identify putative mRNA targets of differentially expressed miRNAs, resulting in 9,369 target genes. These algorithms assess target seed sequence pairing, site count, conservation, and site context scores to reliably predict high-confidence targets. Among these 9,369 target genes, 368 overlapped with upregulated DEGs, and 133 overlapped with downregulated DEGs. Upon examining the negative correlation between DEGs and miRNAs, we identified 239 upregulated genes and 119 downregulated genes (Fig. 5A, Supplementary Fig. 3A-B). To determine whether these overlaps exceeded those expected by chance, we performed two-sided Fisher’s exact tests using the 26,566 genes included in the differential-expression and miRNA-target analyses as the background universe. Upregulated genes were significantly enriched among the corresponding predicted miRNA targets, with 239 genes observed compared with 205.6 expected by chance (fold enrichment = 1.16; odds ratio = 1.23; BH-adjusted *P* = 0.008), whereas no significant enrichment was observed for downregulated genes. To elucidate the molecular pathways affected by the 22q11.2 deletion, GO analysis of these DEGs was conducted (Supplementary Fig. 3C-D). Results revealed that these 239 upregulated DEGs in synaptosomes of 22q11DS organoids are enriched in biological pathways related to nervous system development, PI3K-Akt signaling and MAPK cascades, cytokine response, cytokine production, inflammatory response pathways, and immune system processes (Supplementary Fig. 3C). In contrast, the 119 downregulated DEGs are associated with 22q11.2 copy number variation syndrome, chemical synaptic transmission, trans-synaptic signaling, regulation of neuronal synaptic plasticity, and activation of AMPA receptors (Supplementary Fig. 3D), which is consistent with both disruption in homeostatic plasticity in 2D cultures and decrease of these pathways in 3D organoids.

**Figure 5.**
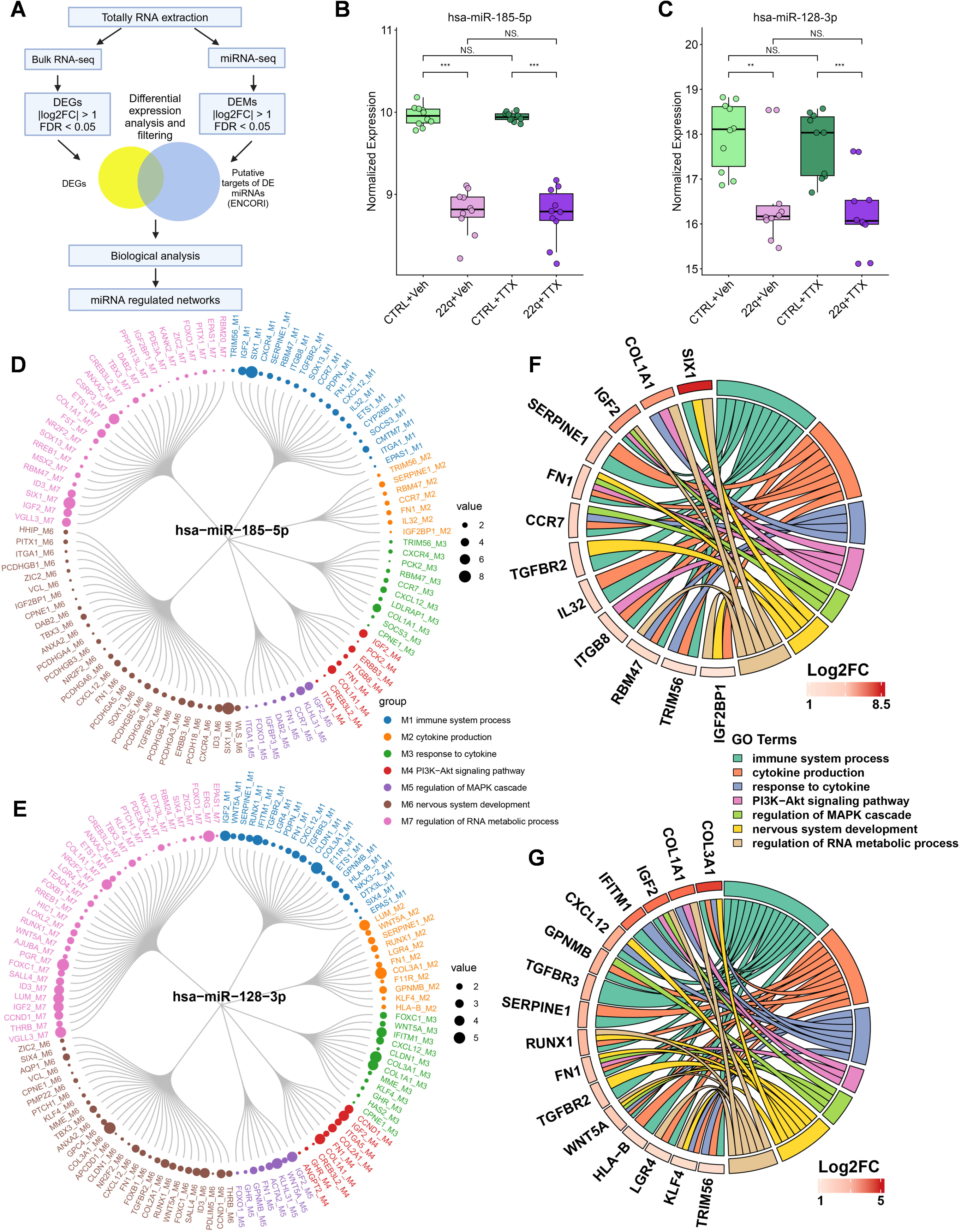
Integrated analysis of miRNA and mRNA profiles in 22q11.2DS synaptosomes. (**A**) Schematic of integrated analyses of miRNA and mRNA profiles from synaptosomes isolated from day 98 cortical organoids derived from healthy control iPSC lines (004, 006, 007, 008, and C19), 22q11DS patient iPSC lines (22q001, 22q002, 22q003, and 22q005), and the CRISPR-engineered C19 isogenic 22q11.2 deletion line (1F3), with two replicates per sample. (**B**) Normalized expression levels of hsa-miR-185-5p. (**C**) Normalized expression levels of hsa-miR-128-3p. (**D**) The key miRNA hsa-miR-185-5p regulates DEGs associated with diverse GO terms in 22q11DS synaptosomes. (**E**) The key miRNA hsa-miR-128-3p regulates DEGs associated with diverse GO terms in 22q11DS synaptosomes. (**F**) Circos plot shows key genes regulated by hsa-miR-185-5p, enriched in immune system processes, cytokine production, and nervous system development. (**G**) Circos plot shows key genes regulated by hsa-miR-128-3p, enriched in immune system processes, cytokine production, and nervous system development. For panels F and G, all displayed genes were significantly differentially expressed at an FDR-adjusted *P* < 0.05; log2 fold-change values are shown to indicate the direction and magnitude of expression changes. GO: gene ontology. DEG: differentially expressed gene. DEM: differentially expressed miRNA.

Notably, the miRNAs hsa-miR-185-5p (Fig. 5B), hsa-miR-185-3p, hsa-miR-1286, and hsa-miR-1306-5p, all located within the 22q11.2 deletion region, were significantly downregulated in 22q11DS synaptosomes compared to controls (Supplementary Fig. 4A-D). Interestingly, hsa-miR-128-3p, which is not located within the deleted region, was also significantly downregulated in 22q11DS synaptosomes (Fig. 5C). To more precisely map the miRNA-regulated pathways that may contribute to 22q11DS pathogenesis, we constructed integrative networks of the 22q11.2 deletion-modulated miRNAs and their target mRNAs (Fig. 5D-E). Intriguingly, upregulated DEGs in 22q11DS synaptosomes are mainly involved in immune system process, cytokine production, response to cytokine, nervous system development, and regulation of RNA metabolic process, many of which are regulated by hsa-miR-185-5p (Fig. 5D) and hsa-miR-128-3p (Fig. 5E). To identify hub genes enriched in overlapping GO terms in 22q11DS synaptosomes, we performed cross-examination (Fig. 5F-G). The Circos plot showed that several key differentially expressed mRNAs—such as *IGF2*, *TRIM56*, *SIX1*, *SERPINE1*, *RBM47*, *ITGB8*, *TGFBR2*, *CCR7*, *FN1*, *IL32*, *IGF2BP1*, and *COL1A1*—were upregulated following downregulation of hsa-miR-185-5p, due to the 22q11.2 deletion (Fig. 5F). In parallel, the network displayed *IGF2*, *TRIM56*, *WNT5A*, *RUNX1*, *IFITM1*, *CXCL12*, *LGR4*, *FN1*, *COL1A1*, *COL3A1*, *HLA-B*, *GPNMB*, *KLF4*, *TGFBR2*, *TGFBR3*, and *SERPINE1*, which were also upregulated as a result of hsa-miR-128-3p downregulation (Fig. 5G). Collectively, these integrative networks highlight a set of dysregulated miRNAs and their downstream targets perturbed by the 22q11.2 deletion, offering mechanistic insights into synaptic dysfunction and immune-related pathways in 22q11DS synapses. Based on their significant downregulation and predicted regulation of multiple DEGs associated with immune and neuronal pathways, we prioritized hsa-miR-185-5p and hsa-miR-128-3p for further investigation.

### hsa-miR-185-5p and hsa-miR-128-3p are key drives for synaptic transcriptome of 22q11DS

Homeostatic synaptic plasticity is driven by synaptic transcriptome remodeling. Strikingly, in control synaptosomes, TTX treatment resulted in the upregulation of 214 genes and downregulation of 177 genes (Fig. 6A; Supplementary Table 5). In contrast, synaptosomes from 22q11DS organoids showed minimal transcriptional changes in response to TTX (Fig. 6B; Supplementary Table 6), further indicating that the 22q11.2 deletion impairs homeostatic synaptic plasticity by altering the synaptic transcriptome at the molecular level. To identify the key driver miRNAs mediating the synaptic deficits in 22q11DS, we used predictive algorithms available in ENCORI^49^ to predict potentially downregulated miRNAs based on the 214 mRNAs upregulated in control synaptosomes following TTX treatment, identifying 216 candidate miRNAs. We then compared these predicted miRNAs with the significantly downregulated DEMs identified by miRNA-seq in 22q11DS versus control synaptosomes under vehicle (37 miRNAs) and TTX (23 miRNAs) conditions. UpSet plot analysis of these three miRNA sets identified hsa-miR-185-5p and hsa-miR-128-3p as overlapping candidates (Fig. 6C). These findings support the role of hsa-miR-185-5p and hsa-miR-128-3p as key mediators of synaptic dysfunction in 22q11DS, potentially linking miRNA dysregulation to aberrant synaptic transmission and plasticity.

**Figure 6.**
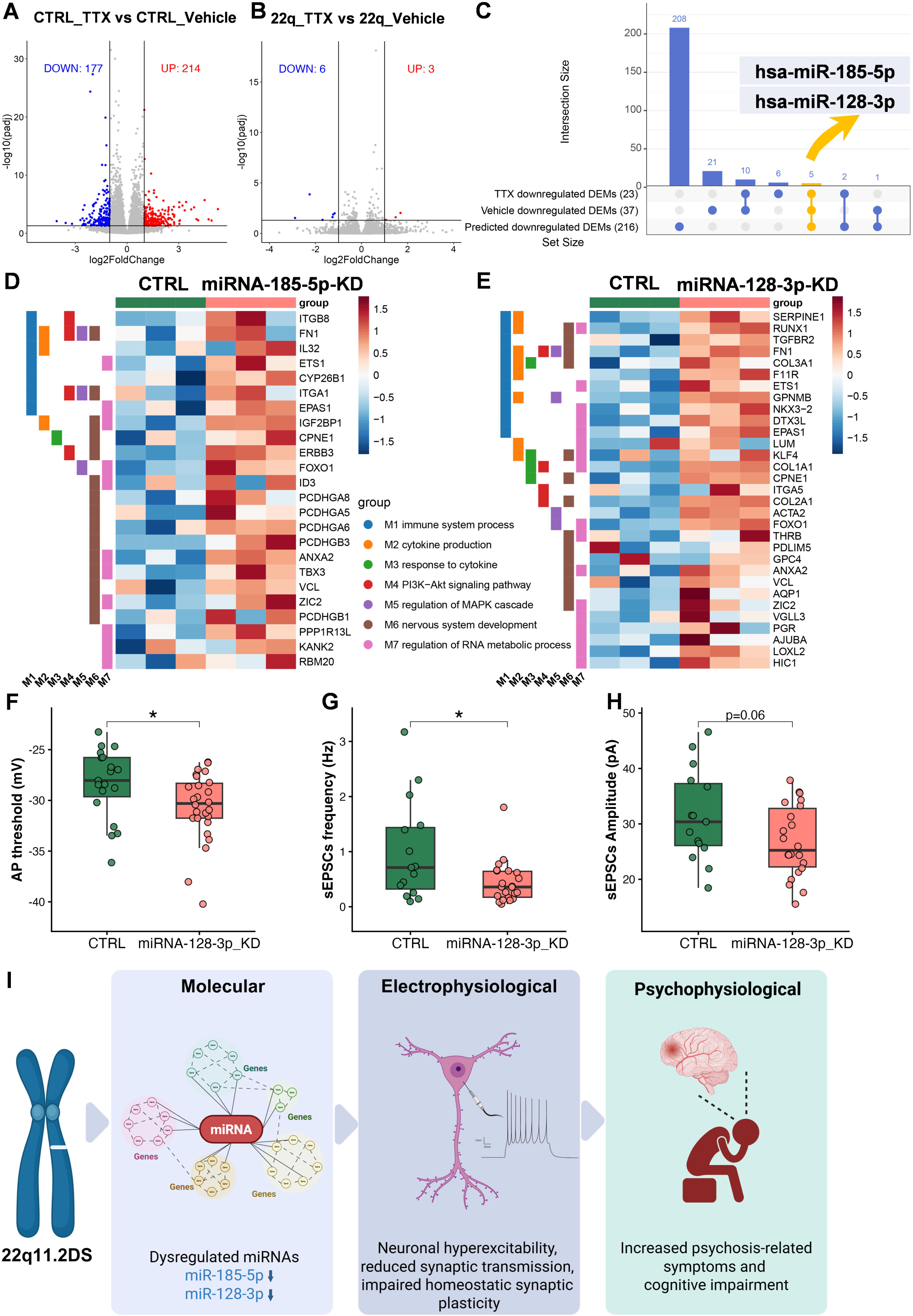
Validation of dysregulated miRNAs in 22q11DS neurons. (**A**) Volcano plot shows DEGs in synaptosomes isolated from day 98 control cortical organoids (004, 006, 007, 008, and C19), with two replicates per sample, following 24-hour treatment with either vehicle (DMSO) or TTX. Red and blue dots represent 214 upregulated and 177 downregulated genes, respectively (adjusted *P*-value < 0.05; absolute log2fold-change > 1.0). Grey dots indicate genes without significant differential expression. (**B**) Volcano plot shows DEGs in synaptosomes isolated from day 98 22q11DS cortical organoids (22q001, 22q002, 22q003, 22q005, and the CRISPR-engineered C19 isogenic 22q11.2 deletion line 1F3), with two replicates per sample, following 24-hour treatment with either vehicle (DMSO) or TTX. Red and blue dots represent 3 upregulated and 6 downregulated genes, respectively (adjusted *P*-value < 0.05; absolute log2fold-change > 1.0). Grey dots indicate genes without significant differential expression. (**C**) UpSet plot showing the intersections among 216 candidate downregulated miRNAs predicted from the 214 mRNAs upregulated in control synaptosomes following TTX treatment, 37 significantly downregulated DEMs identified in 22q11DS versus control synaptosomes under vehicle conditions, and 23 significantly downregulated DEMs identified in 22q11DS versus control synaptosomes under TTX conditions (adjusted *P* < 0.05, absolute log2 fold-change > 1). (**D**) Heatmap of hsa-miR-185-5p regulated genes associated with diverse GO terms in control and hsa-miR-185-5p knockdown (miR185-KD) neurons, with three replicates per group. (**E**) Heatmap of hsa-miR-128-3p–regulated genes associated with diverse GO terms in control and hsa-miR-128-3p knockdown (miR128-KD) neurons, with three replicates per group. (**F-H**) Electrophysiological properties of cortical neurons from 6-week-old healthy controls and hsa-miR-128-3p KD were evaluated using patch-clamp recordings. Shown are quantifications of action potential threshold (F; data are presented as mean ± s.e.m.; n = 19 CTRL, n = 26 miR128-KD; \**P* < 0.05, student’s t-test), and the frequency (G) and amplitude (H) of sEPSCs (Data are presented as mean ± s.e.m.; n = 15 CTRL, 22 miR128-KD; \**P* < 0.05, Student’s t-test). (**I**) The diagram illustrates the molecular, electrophysiological, and psychosis-related clinical and psychophysiological mechanisms underlying 22q11.2DS. Dysregulated expression of specific miRNAs, including the downregulation of hsa-miR-185-5p and hsa-miR-128-3p, in 22q11DS neurons leads to neuronal hyperexcitability, aberrant glutamatergic synaptic transmission, and disrupted homeostatic synaptic plasticity, which collectively contribute to psychosis-related clinical and psychophysiological phenotypes. DEGs: differentially expressed genes; DEMs: differentially expressed miRNAs; sEPSCs: spontaneous excitatory postsynaptic currents.

To further investigate the causal role of hsa-miR-185-5p and hsa-miR-128-3p in the increased inflammation and decreased synaptic transmission observed in 22q11DS neurons, we knocked down hsa-miR-185-5p or hsa-miR-128-3p in a control line (004 cell line), respectively. We validated efficient knockdown of hsa-miR-185-5p in cortical neurons by qPCR, which showed a significant reduction of miR-185-5p levels in knockdown (miR185-KD) neurons relative to controls (Supplementary Fig. 5A). Hierarchical clustering of bulk RNA-seq data showed distinct separation between CTRL and miR185-KD samples (Supplementary Fig. 5B). GO analyses identified the upregulation of 898 genes (Supplementary Fig. 5C; Supplementary Table 7) in neurons with hsa-miR-185-5p knockdown associated with immune system process, immune response, cytokine production, and chemokine production (Supplementary Fig. 5D); while 355 downregulation genes were associated with trans-synaptic signaling, chemical synaptic transmission, postsynaptic membrane potential, and nervous system process (Supplementary Fig. 5E). Gene set variation analysis (GSVA) further confirmed increased enrichment of immune-related gene sets (Supplementary Fig. 5F-I), including immune response (Supplementary Fig. 5F), immune system development (Supplementary Fig. 5G), positive regulation of cytokine production (Supplementary Fig. 5H), and chemokine production (Supplementary Fig. 5I); while decreased expression of learning (Supplementary Fig. 5J), memory (Supplementary Fig. 5K), cognition (Supplementary Fig. 5L), positive regulation of excitatory postsynaptic potential (Supplementary Fig. 5M), regulation of glutamatergic synaptic transmission (Supplementary Fig. 5N), and regulation of synaptic plasticity (Supplementary Fig. 5O). We next examined whether miR-185-5p knockdown recapitulated the expression changes of the miR-185-5p target genes identified through our integrative analysis (Fig. 5D). A set of these target genes, including*ITGB8*, *FN1*, *IL32*, *ETS1*, *CYP26B1*, *ITGA1*, *EPAS1*, *IGF2BP1*, *CPNE1*, *ERBB3*, *FOXO1*, *ID3*, *PCDHGA8*, *PCDHGA5*, *PCDHGA6*, *PCDHGB3*, *ANXA2*, *TBX3*, *VCL*, *ZIC2*, *PCDHGB1*, *PPP1R13L*, *KANK2*, and *RBM20* were upregulated in miR185-KD neurons compared with controls (Fig. 6D), consistent with their dysregulation in 22q11DS neurons.

Similarly, knockdown of hsa-miR-128-3p in cortical neurons resulted in comparable transcriptomic alterations (Supplementary Fig. 6A-O). qPCR confirmed effective reduction of miR-128-3p levels in knockdown (miR128-KD) neurons (Supplementary Fig. 6A). Further transcriptomic analyses revealed upregulation of immune-related genes and downregulation of synaptic and cognitive function-associated pathways (Supplementary Fig. 6B-E; Supplementary Table 8). GSVA showed consistent enrichment patterns with those observed in miR-128-3p knockdown neurons, including enhanced immune signatures (Supplementary Fig. 6F-I) and reduced signatures related to cognition and synaptic function (Supplementary Fig. 6J-O). Importantly, a set of hsa-miR-128-3p target genes identified as upregulated DEGs in 22q11DS synaptosomes through integrative analysis (Fig. 5E) were also increased in miR128-KD neurons, including *SERPINE1*, *RUNX1*, *TGFBR2*, *FN1*, *COL3A1*, *F11R*, *ETS1*, *GPNMB*, *NKX3-2*, *DTX3L*, *EPAS1*, *LUM*, *KLF4*, *COL1A1*, *CPNE1*, *ITGA5*, *COL2A1*, *ACTA2*, *FOXO1*, *THRB*, *PDLIM5*, *GPC4*, *ANXA2*, *VCL*, *AQP1*, *ZIC2*, *VGLL3*, *PGR*, *AJUBA*, *LOXL2*, and *HIC1* (Fig. 6E).

Furthermore, electrophysiological recordings revealed that miR128-KD neurons mirror the functional deficits observed in 22q11DS neurons. Patch-clamp recording analyses showed a significantly decreased AP threshold in miR128-KD neurons compared to controls (Fig. 6F), indicating enhanced neuronal excitability. Moreover, sEPSCs exhibited markedly reduced frequency (Fig. 6G) and a trend toward decreased amplitude in miR128-KD neurons (Fig. 6H; *P*= 0.06), consistent with impaired glutamatergic transmission in 22q11DS neurons. Collectively, these results suggest hsa-miR-185-5p and hsa-miR-128-3p as key regulators of synaptic transcriptome remodeling in 22q11DS, potentially acting in concert to drive immune dysregulation and synaptic dysfunction in 22q11DS cortical neurons.

## DISCUSSION

In this study, we leveraged a comprehensive multimodal platform, including extensive cellular, electrophysiological, integrative transcriptomic, psychosis-related clinical and psychophysiological analyses, to elucidate the neuronal abnormalities underlying 22q11DS. Our results demonstrate that the 22q11.2 deletion promotes neuronal hyperexcitability, disrupts glutamatergic synaptic transmission and homeostatic synaptic plasticity. Notably, we identify dysregulated miRNAs as key mechanistic mediators linking the genomic deletion to synaptic dysfunction. Together, these findings illustrate the molecular, electrophysiological, and psychosis-related clinical and psychophysiological mechanisms underlying 22q11.2DS (Fig. 6I). hsa-miR-185-5p and hsa-miR-128-3p act as key drivers of synaptic transcriptome remodeling in 22q11DS, synergistically mediating immune and synaptic dysfunction in cortical neurons. Collectively, these alterations may underlie psychosis-related clinical and psychophysiological phenotypes (Fig. 6I), offering new insight into the pathophysiology of 22q11DS and informing future therapeutic approaches for 22q11DS and SCZ.

The observed dissociation between intrinsic and synaptic properties in 22q11DS neurons, characterized by a lowered AP threshold and increased fAHP alongside a decreased frequency of sEPSCs, points to a profound state of circuit-level dysregulation. This profile suggests a compensatory, yet ultimately maladaptive, homeostatic response to reduced excitatory drive^64^. In models of neurodevelopmental disorders such as ASD and SCZ, the primary loss of functional synapses or excessive pruning (sEPSC reduction) appears to trigger a “scaling up” of intrinsic excitability as the neuron attempts to maintain a target firing rate. By lowering the AP threshold, the neuron increases its sensitivity to the remaining sparse inputs; concurrently, the upregulation of BK channels, evidenced by the increased fAHP, facilitates high-frequency repetitive firing by accelerating the recovery of voltage-gated sodium channels from inactivation^65^. However, this intrinsic “gain” increase likely compromises the signal-to-noise ratio of the circuit. While the neuron becomes easier to fire, it does so in a manner that is increasingly decoupled from organized network activity, potentially explaining the sensory hypersensitivity and cognitive fragmentation hallmark to these conditions^66^. This “hyper-responsive yet disconnected” state mirrors findings in Fmr1 knockout models and the prefrontal cortical deficits observed in SCZ, where the metabolic cost of maintaining such high intrinsic readiness may eventually contribute to cellular stress or excitotoxicity^67,68^. Thus, our findings suggest a complex interaction between structural synaptic loss and compensatory membrane adaptations. Consistent with our results, previous iPSC-based studies have also reported that neurons derived from individuals with 22q11DS exhibit reduced network activity and aberrant calcium signaling^19^. Nehme et al. reported that the 22q11.2 deletion exerts widespread trans effects on gene expression in human iPSC-derived excitatory neurons, leading to dysregulation of presynaptic factors, proteasome components, and JUN/FOS transcriptional regulators^20^. Their integrative network analysis revealed that these perturbed gene modules converge with known schizophrenia risk pathways, highlighting presynaptic and transcriptional dysfunction as core mechanisms through which 22q11.2 haploinsufficiency confers psychiatric risk^20^. Recent organoid studies reinforce a neurodevelopmental basis for schizophrenia-associated phenotypes in 22q11DS^21^. Cortical organoids derived from affected individuals show delayed neuronal maturation, increased progenitor proliferation, and fewer mature excitatory neurons. Single-cell analyses reveal disrupted differentiation programs, impaired neurite development, and blunted glutamate-evoked Ca²⁺ signaling^21^. Notably, whereas the organoid study reported reduced neurite length and complexity, our 22q11DS NGN2-induced neurons exhibited increased neurite complexity. This apparent discrepancy may reflect differences in the differentiation paradigms, cellular context, and developmental stage between 2D NGN2-induced neurons and 3D cortical organoids. Together, these findings highlight early cortical maturation defects and synaptic pathway disruptions as cellular mechanisms through which the 22q11.2 deletion elevates schizophrenia risk. Despite the clinical heterogeneity observed among individuals with 22q11DS, these cellular phenotypes were robust and reproducible across different subjects and cell lines, suggesting a convergence at the molecular and cellular levels.

To our knowledge, this study provides the first evidence that 22q11DS neurons exhibit profoundly impaired homeostatic synaptic plasticity, a critical negative feedback mechanism that stabilizes neuronal activity in response to perturbations^54,55,69,70^. Upon chronic activity blockade, control neurons showed the expected homeostatic increase in mEPSC amplitude, whereas 22q11DS neurons failed to mount a compensatory response (Fig. 3D-H). Transcriptomic analyses of neurons and synaptosomes of cortical organoids revealed broad dysregulation of synapse-related pathways. Downregulated genes were enriched for glutamatergic synaptic transmission, regulation of neuronal synaptic plasticity, and AMPA receptor activity (Supplementary Fig. 3D), all of which align with the electrophysiological impairments observed. Conversely, genes associated with inflammation, cytokine signaling, and immune activation were significantly upregulated. These findings indicate that synaptic dysfunction in 22q11DS is accompanied by aberrant neuroinflammation, which may further destabilize neural functions. Immune dysregulation is a well-recognized feature of 22q11DS, with abnormalities in immune cell development and function, increased susceptibility to autoimmunity, and altered inflammatory profiles reported in affected individuals^71–73^. Moreover, elevated pro-inflammatory markers have been associated with psychosis and cognitive deficits in 22q11DS^74^, suggesting a potential link between immune dysregulation and neuropsychiatric manifestations. Emerging evidence also implicates blood-brain barrier dysfunction and altered neuroimmune interactions in 22q11DS^75,76^, further supporting a potential contribution of immune-related mechanisms to central nervous system dysfunction. In this context, the recurrent enrichment of immune-, cytokine-, and chemokine-related pathways in our neuronal and synaptosome transcriptomic analyses suggests that immune-related molecular dysregulation may extend to neuronal compartments and intersect with synaptic dysfunction. Notably, we also observed enrichment of immune-and inflammatory-related pathways following miRNA perturbation, raising the possibility that altered miRNA-mediated gene regulation contributes to these transcriptional signatures. Given the established role of DGCR8 haploinsufficiency and impaired miRNA biogenesis in 22q11DS, these findings suggest a potential molecular link between the 22q11.2 deletion, miRNA dysregulation, altered neuronal immune-related signaling, and synaptic dysfunction. Future studies incorporating microglia and other immune components into human iPSC-derived neuronal and organoid models will be important for defining the functional interactions among these processes. GSVA confirmed that 22q11DS synaptosomes fail to activate transcriptomic gene sets associated with synapse-related pathways in response to synaptic silencing (Fig. 4E-H), providing molecular evidence for impaired synaptic homeostasis in 22q11DS. These findings support the notion that impaired synaptic plasticity may be a key contributor to the neuropathology and cognitive dysfunction observed in schizophrenia^11,12^. Postmortem studies of the dorsolateral prefrontal cortex have reported significant reductions in transcripts associated with synaptic plasticity and cytoskeletal remodeling in individuals with schizophrenia^77^. Therefore, delineating disease-associated synaptic deficits and their underlying mechanisms is essential for understanding the pathogenesis of 22q11DS and schizophrenia and for guiding the development of novel therapeutic interventions.

A critical novel insight from our study is the role of dysregulated synaptic miRNAs in mediating the effects of the 22q11.2 deletion. MicroRNAs are increasingly recognized as critical contributors to the pathophysiology of psychiatric disorders, including schizophrenia, bipolar disorder, and depression. These small non-coding RNAs regulate gene expression by binding to target mRNAs, resulting in translational repression or degradation^78^. Their ability to fine-tune gene networks makes them key contributors to brain development, synaptic plasticity, and stress responses—all processes implicated in psychiatric disorders^79–82^. Heterozygous loss of DGCR8 has been shown to recapitulate the excitability and calcium signaling phenotypes of 22q11DS neurons, whereas its overexpression rescues these defects^19^, further implicating impaired miRNA biogenesis as a pathogenic mechanism underlying neuronal dysfunction in 22q11DS. Our integrative mRNA-miRNA network analysis in 22q11DS synaptosomes revealed that downregulated miRNAs, including hsa-miR-185-5p and hsa-miR-128-3p, target genes involved in synaptic signaling and immune system process. These miRNAs are predicted to regulate multiple upregulated genes in 22q11DS synaptosomes, including *COL1A1*, *IGF2*, *FN1*, and *TGFBR2*, all of which have been implicated in neurodevelopmental disorders^83–87^. MiR-185-5p, located within the 22q11.2 deletion region, is the only miRNA consistently downregulated across three mouse model studies^88–90^ and two human induced pluripotent stem cell models of the 22q11.2 deletion^91,92^. Its downregulation represents a shared pathogenic mechanism in both 22q11.2DS–associated and idiopathic schizophrenia^93^. These findings suggest that alterations in gene expression mediated by miR-185-5p may be a mechanistic connection between SCZ and 22q11.2DS. Importantly, previous work further supports the translational relevance of miR-185 dysregulation in 22q11.2DS. Thakur et al. demonstrated reduced miR-185 expression and increased expression of its downstream target EMC10 in human iPSC-derived neurons from individuals with 22q11.2DS. Reducing EMC10 expression ameliorated neuronal phenotypes in patient-derived neurons, while antisense oligonucleotide-mediated reduction of Emc10 improved cognitive deficits in a 22q11.2 deletion mouse model, highlighting the therapeutic potential of targeting downstream pathways affected by miR-185 dysregulation^94^.

Notably, we observed reduced miR-128-3p expression in 22q11DS synaptosomes. Hsa-miR-128-3p is not located within the deleted region, suggesting that downstream regulatory cascades may propagate the impact of the deletion beyond its direct genomic footprint. MiR-128-3p plays a critical role in mammalian brain development, being prominently expressed in differentiating neuronal cells and characterized by its highly tissue-specific expression and regulation across developmental stages^95^. It is highly expressed in the human brain^96^ (Supplementary Fig. 7) and functions from the early regulation of neural progenitor cells to guiding the formation of complex neuronal circuits. Among its targets, miR-128-3p regulates the expression of a broad set of genes, including those encoding for ion channels and transporters, which are critical for maintaining the electrical activity of neurons that contribute to neuronal excitability and neurotransmitter-driven motor activity^97^. Collectively, our findings implicate miRNA dysregulation as a convergent mechanism connecting 22q11DS with psychiatric disorders, including schizophrenia, and underscore miRNAs as potential therapeutic targets.

In summary, our study provides a detailed mechanistic framework linking the 22q11.2 deletion to synaptic dysfunction via disrupted glutamatergic synaptic transmission, impaired homeostatic plasticity, and miRNA-mediated transcriptional dysregulation. These findings advance our understanding of the molecular mechanism of neuropsychiatric symptoms in 22q11DS and offer new targets for therapeutic intervention. The pronounced symptom heterogeneity in 22q11DS poses major challenges for elucidating the underlying mechanisms and for developing effective therapeutic interventions. Future studies should extend these findings to larger and more diverse patient cohorts. Incorporating longitudinal assessments and in vivo models will further elucidate the developmental trajectories of these neuronal abnormalities and their contribution to psychosis onset.

## ACKNOWLEDGEMENTS

We thank the members of the Wen lab for helpful discussions. This work was supported in part by NIH grants (R01MH139822, R01MH121102, R01AG065611, and R21MH132012 to ZW; R01MH117315 to JFC and ED; UL1TR002378 TL1TR002382 to DP). The NARSAD Young Investigator Grant from the Brain & Behavior Research Foundation (BBRF) to WN (James John Karatheodoris Memorial Investigator, Grant Number: 33517) and the TSC Alliance Research Grant to WN (Wong Family Foundation Research Award, Crossref DOI: 1340014); Department of Psychiatry and Behavioral Sciences within the Emory University School of Medicine general support funds to OYO.

## AUTHOR CONTRIBUTIONS

WN led and was involved in all aspects of the project. WN performed cell culture and cellular experiments and wrote the initial manuscript draft. XR, SY and WN performed the transcriptomic analyses. JG led and performed the patch-clamp recordings with assistance from XL, CM, EG, SYJ and EX assisted with cell maintenance and sample collection. DP, OYO, BP, EFW, JFC, and ED conducted clinical assessments. ED secured IRB approval, obtained informed consent from the families, and collected blood samples from the subjects. SI contributed to the collection of peripheral blood samples. RP, GJB, and JGM provided the isogenic cell lines. ZW, ED, ZQ, JFC, YF, and CEB supervised the project. ZW, ED, JG, and WN designed the project and finalized the manuscript.

## ADDITIONAL INFORMATION

## Study Approval

All studies adhered to institutional IRB protocols approved by the Emory University School of Medicine.

## Data and materials availability

All data are available in the main text or the supplementary materials. We have deposited the RNA-seq and smRNA-seq data into the Gene Expression Omnibus (NCBI). The accession numbers are GSE294898 and GSE304650.

## Code availability

Codes are available at https://github.com/ranxm2/22q_mutilomics.

