## Supplementary Information for "Synaptic microRNA dysregulation orchestrates aberrant glutamatergic transmission in a human neuronal model of 22q11.2 deletion syndrome"

[1]Department of Psychiatry and Behavioral Sciences, Emory University School of Medicine, Atlanta, Georgia, USA.
[2]Department of Biostatistics and Bioinformatics, Rollins School of Public Health, Emory University, Atlanta, Georgia, USA.
[3]Department of Computer Science, Emory University, Atlanta, Georgia, USA.
[4]Department of Psychology, Emory University, Atlanta, Georgia, USA.
[5]College of Arts and Sciences, Emory University, Atlanta, Georgia, USA.
[6]Atlanta Veterans Administration Health Care System, Decatur, Georgia, USA.
[7]School of Neuroscience, Fralin Biomedical Research Institute at VTC, Virginia Tech, Roanoke, Virginia, USA.
[8]Decatur High School, Decatur, Georgia, USA.
[9]Department of Cell Biology, Emory University School of Medicine, Atlanta, Georgia, USA.
[10]Department of Psychiatry, Rutgers University, New Brunswick, New Jersey, USA.
[11]Department of Epidemiology, Rollins School of Public Health, Emory University, Atlanta, Georgia, USA.
[12]Department of Pharmacology and Chemical Biology, Emory University School of Medicine, Atlanta, Georgia, USA.
[13]Department of Psychiatry and Biobehavioral Sciences, Semel Institute for Neuroscience and Human Behavior, University of California, Los Angeles, California, USA
[14]Department of Psychology, University of California, Los Angeles, California, USA
[15]Department of Human Genetics, Emory University School of Medicine, Atlanta, Georgia, USA.
[16]Department of Neurology, Emory University School of Medicine, Atlanta, Georgia, USA.
[17]These authors contributed equally.

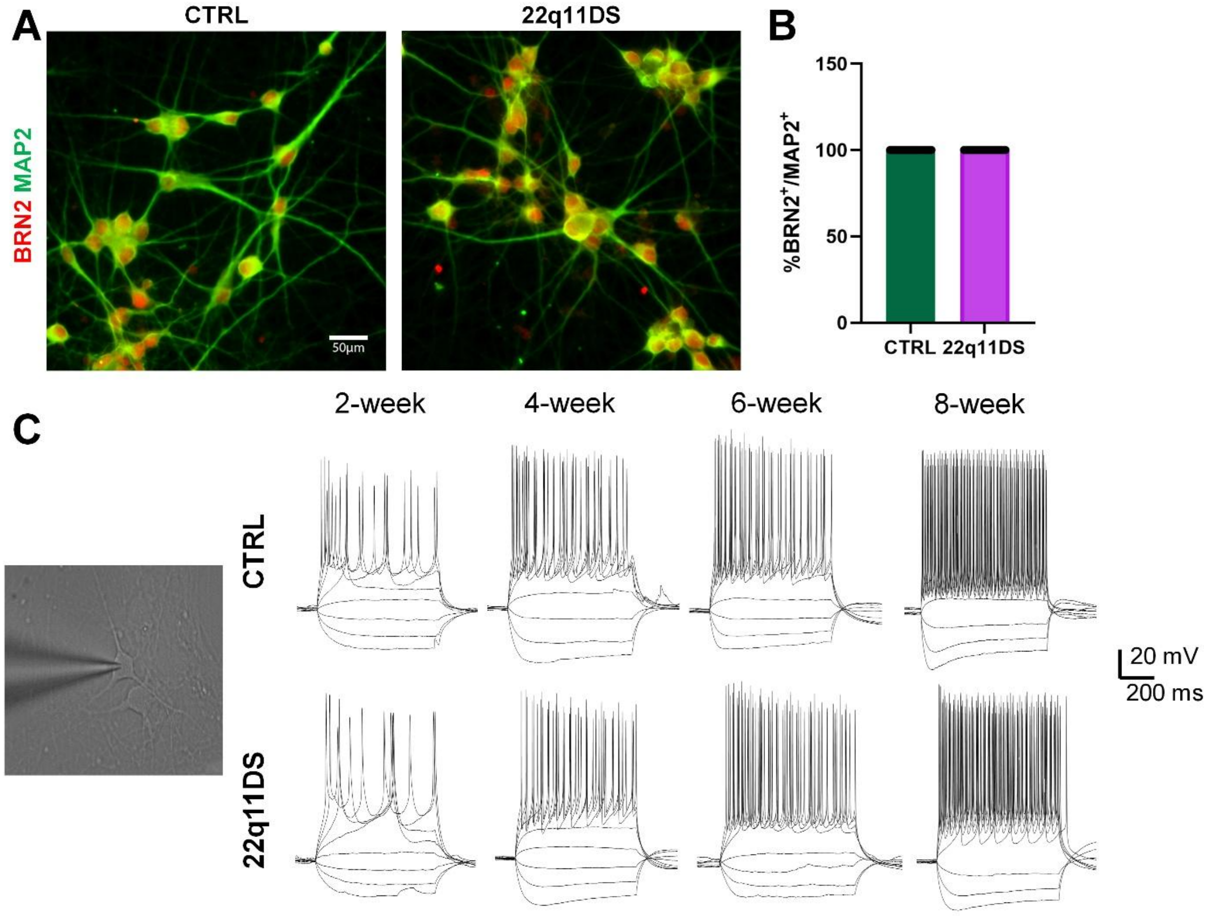


**Supplementary Figure 1. Electrophysiological maturation in 22q11DS cortical neurons.** (**A**) Sample images of hiPSC-derived cortical neurons stained for BRN2 (red) and MAP2 (green) in CTRL and 22q11DS groups. Scale bars, 50 µm. (**B**) Quantification of BRN2$^+$ expression among MAP2$^+$ neurons derived from control lines (004, 006, 007, and 008) and 22q11DS patient lines (22q001, 22q002, 22q003, and 22q005), with 10-14 neurons analyzed per line (total n = 53 CTRL and 46 22q11DS; Student's t-test). (**C**) Representative whole-cell current clamp recordings from CTRL and 22q11DS neurons at 2, 4, 6, and 8 weeks after differentiation.

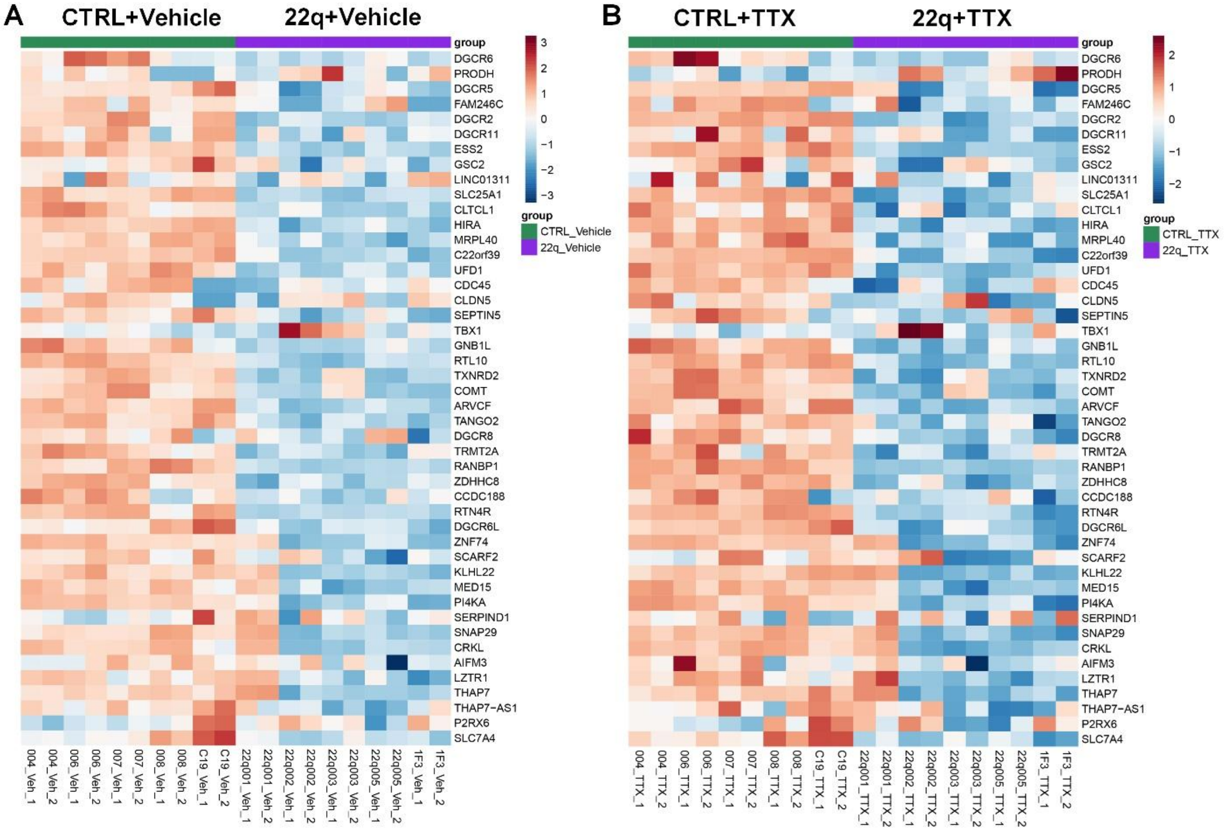


**Supplementary Figure 2. Expression of genes located within the 22q11.2 deletion region in synaptosomes from cortical organoids.** (**A**) Gene expression profiles of synaptosomes isolated from day 98 cortical organoids under vehicle conditions, comparing healthy control lines (004, 006, 007, 008, and C19) and 22q11DS lines (22q001, 22q002, 22q003, 22q005, and the CRISPR-engineered C19 isogenic 22q11.2 deletion line 1F3), with two replicates per sample. (**B**) Gene expression profiles of synaptosomes isolated from day 98 cortical organoids following 24-hour treatment with TTX, comparing healthy control lines (004, 006, 007, 008, and C19) and 22q11DS lines (22q001, 22q002, 22q003, 22q005, and the CRISPR-engineered C19 isogenic 22q11.2 deletion line 1F3), with two replicates per sample. Colors represent normalized gene expression levels. Veh: Vehicle.

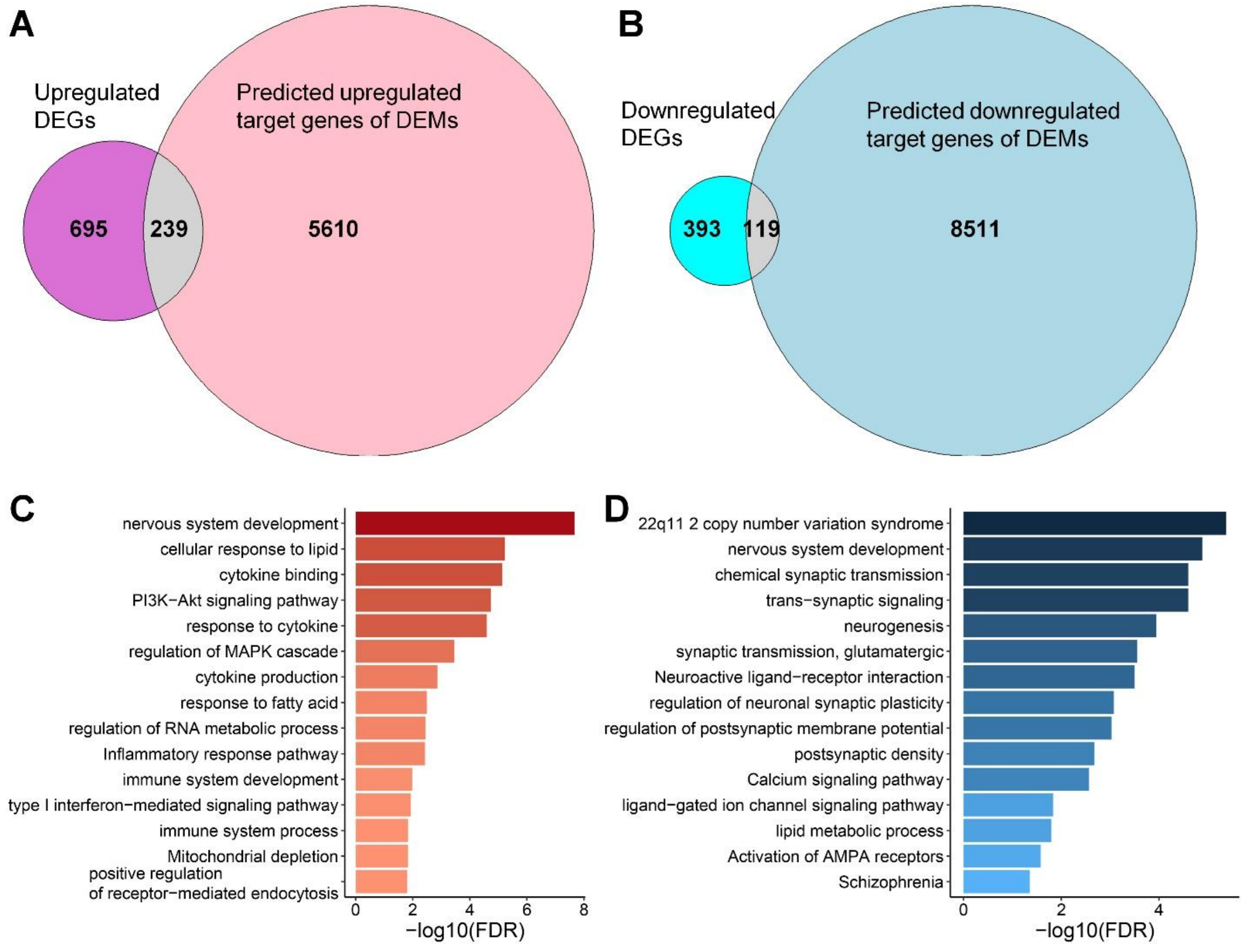


**Supplementary Figure 3. Integrative transcriptome-miRNAome analysis of synaptosomes from control and 22q11DS cortical organoids.** Synaptosomes were isolated from day 98 control cortical organoids (004, 006, 007, 008, and C19), 22q11DS cortical organoids (22q001, 22q002, 22q003, and 22q005), and CRISPR-engineered C19 isogenic 22q11.2 deletion cortical organoids (1F3), with two replicates per sample. (**A-B**) Venn diagrams illustrating the overlap between DEGs and predicted targets of DEMs, including overlap between upregulated DEGs and predicted miRNA targets (A) and overlap between downregulated DEGs and predicted miRNA targets (B) in 22q11DS synaptosomes. (**C-D**) GO analysis of 239 overlap upregulated (C) and 119 downregulated (D) DEGs regulated by miRNAs in 22q11DS synaptosomes (FDR < 0.05). GO: gene ontology. DEG: differentially expressed gene. DEM: differentially expressed miRNA.

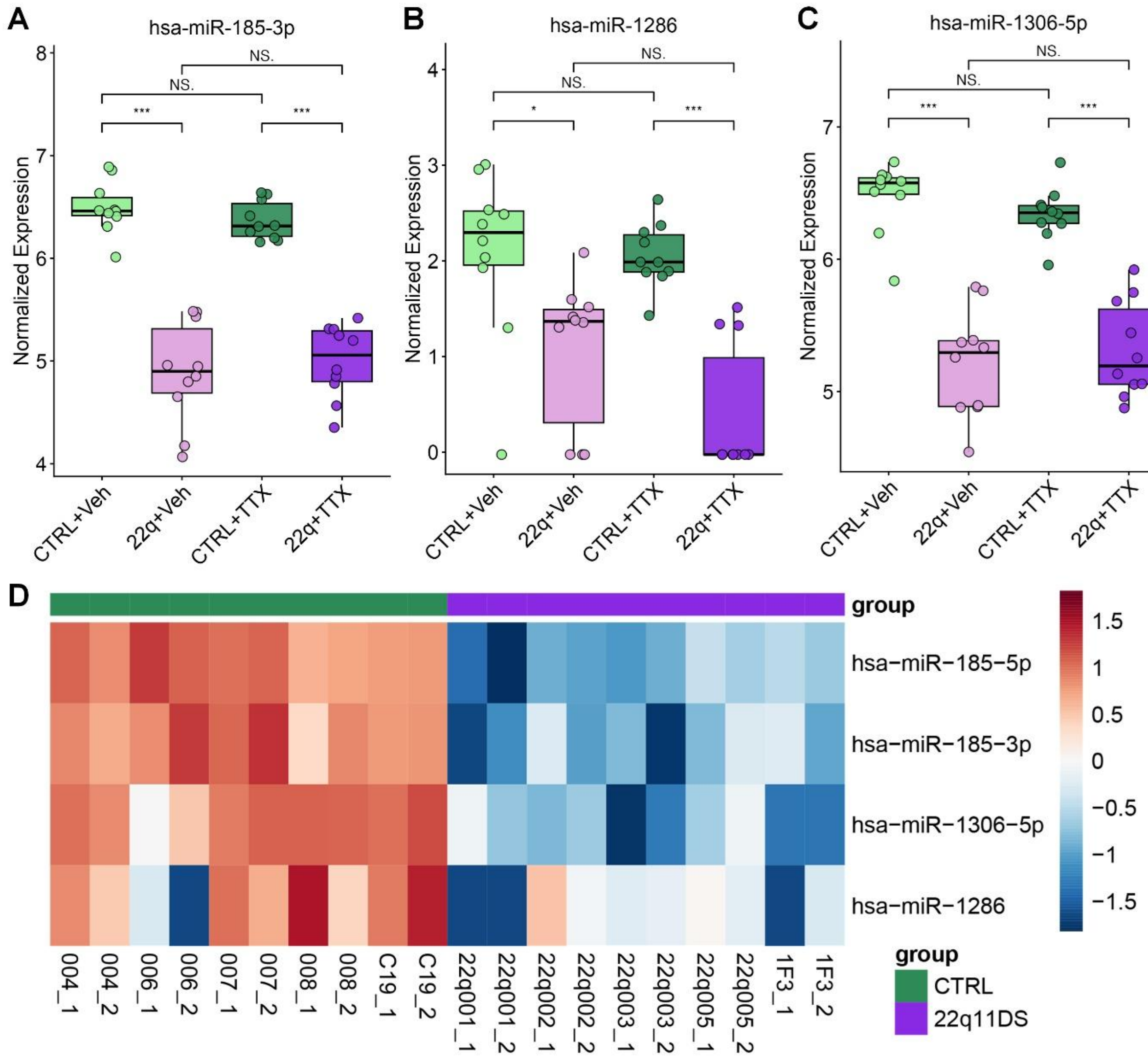


**Supplementary Figure 4. Downregulated miRNAs in synaptosomes from 22q11DS cortical organoids.** Synaptosomes were isolated from day 98 control cortical organoids (004, 006, 007, 008, and C19), 22q11DS cortical organoids (22q001, 22q002, 22q003, and 22q005), and CRISPR-engineered C19 isogenic 22q11.2 deletion cortical organoids (1F3) following 24-hour treatment with vehicle or TTX, with two replicates per sample. (**A-C**) Shown are normalized expression levels of hsa-miR-185-3p (A), hsa-miR-1286 (B), and hsa-miR-1306-5p (C) under vehicle conditions or following 24-hour treatment with TTX. (D) Heatmap shows downregulated miRNAs in synaptosomes from 22q11DS cortical organoids under vehicle conditions across different cell lines. Colors represent normalized miRNA expression levels.

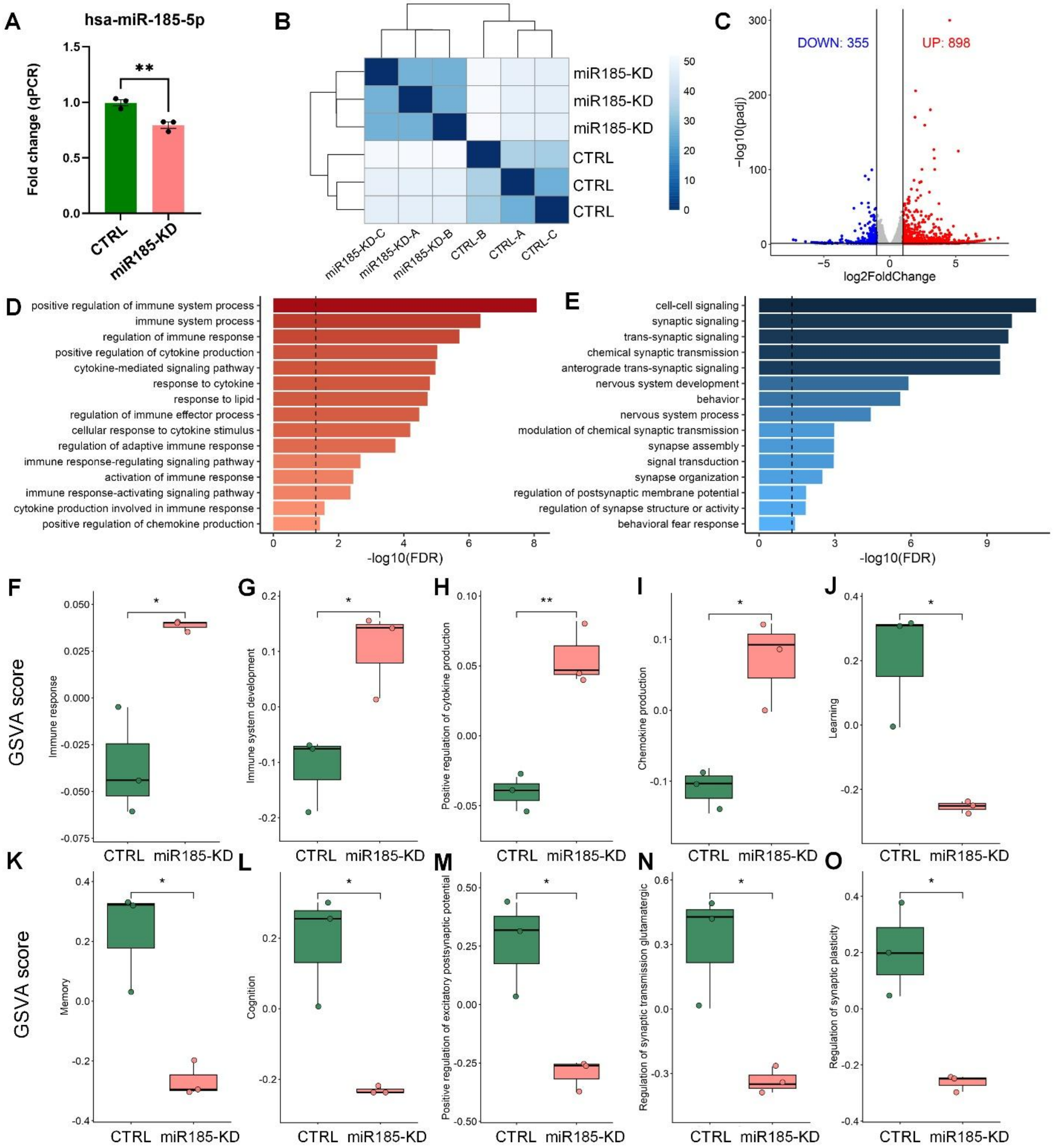


**Supplementary Figure 5. Knockdown of hsa-miR-185-5p increased inflammation, impaired cognitive function, decreased synaptic potential, synaptic transmission, and synaptic plasticity in neurons.** (**A**) RT-qPCR analysis of hsa-miR-185-5p expression in control (CTRL) and hsa-miR-185-5p knockdown (miR185-KD) neurons. Data are presented as mean ± s.e.m. ($n = 3$ for each condition; $^{**}P < 0.01$, unpaired t-test). (**B**) Sample distance heatmap of bulk RNA-seq data from CTRL and miR185-KD neurons. (**C**) Volcano plot shows DEGs in miR185-KD neurons compared to controls. Red and blue dots indicate 898 upregulated genes and 355 downregulated genes, respectively (adjusted $P$-value < 0.05, absolute log2 fold-change > 1.0). Grey dots represent genes with no significant differential expression. (**D-E**) Gene ontology (GO) analyses of upregulated (D) and downregulated (E) DEGs in miR185-KD neurons (FDR < 0.05). (**F-O**) Gene Set Variation Analysis (GSVA) of the transcriptome was performed on control and miR185-KD neurons, including immune response (F), immune system development (G), positive regulation of cytokine production (H), chemokine production (I), learning (J), memory (K), cognition (L), positive regulation of excitatory postsynaptic potential (M), regulation of glutamatergic synaptic transmission (N), and regulation of synaptic plasticity (O). *$P < 0.05$, **$P < 0.01$, Student's t-test. DEGs: differentially expressed genes.

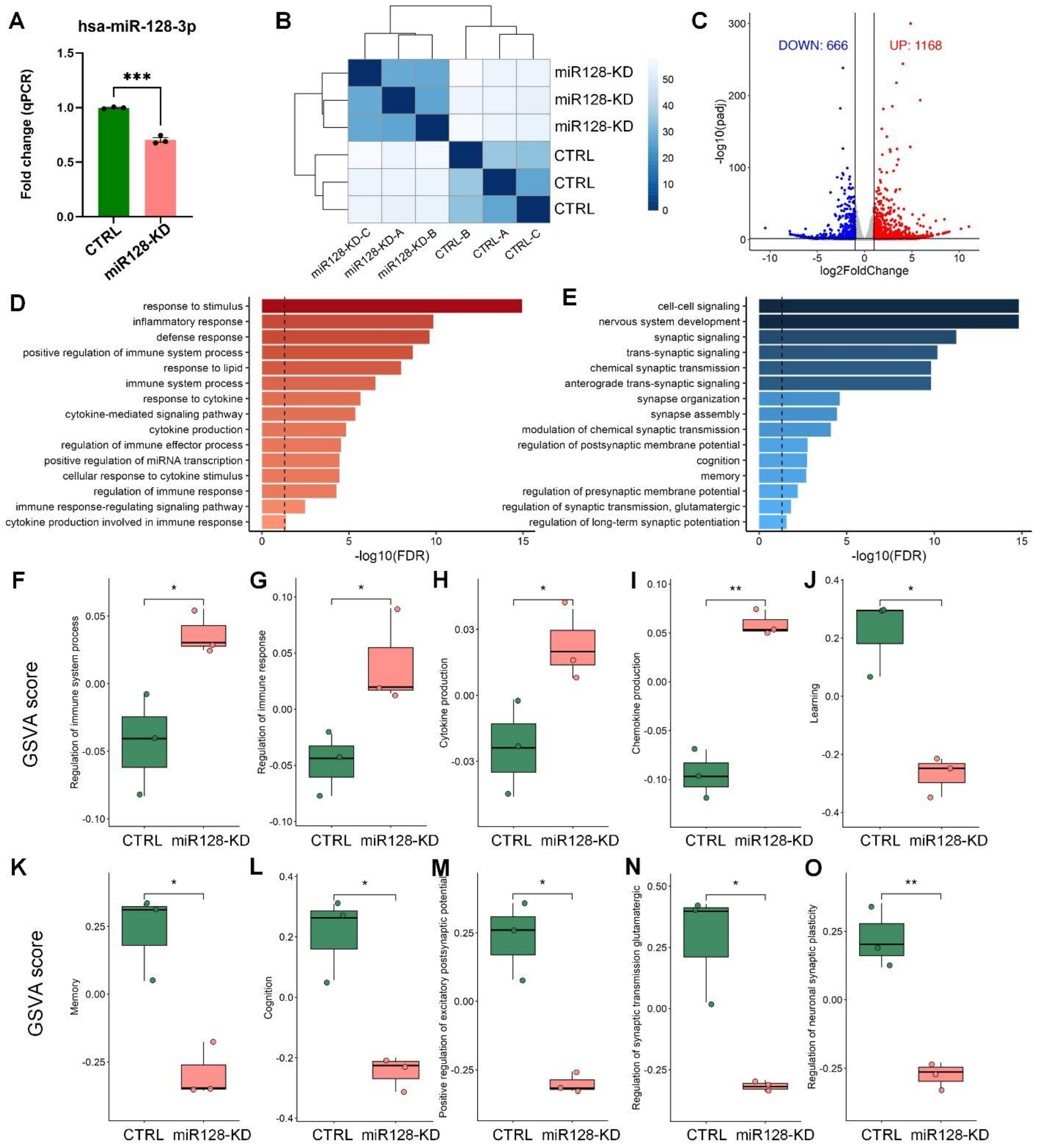


**Supplementary Figure 6. Knockdown of hsa-miR-128-3p increased inflammation, impaired cognitive function, decreased synaptic potential, synaptic transmission, and synaptic plasticity in neurons.** (**A**) RT-qPCR analysis of hsa-miR-128-3p expression in control (CTRL) and hsa-miR-128-3p knockdown (miR128-KD) neurons. Data are presented as mean ± s.e.m. ($n = 3$ for each condition; ****P* < 0.001, unpaired t-test). (**B**) Sample distance heatmap of bulk RNA-seq data from CTRL and miR128-KD neurons. (**C**) Volcano plot shows DEGs in miR128-KD neurons compared to controls. Red and blue dots indicate 1168 upregulated genes and 666 downregulated genes, respectively (adjusted *P*-value < 0.05, absolute log2 fold-change > 1.0).

Grey dots represent genes with no significant differential expression. (**D-E**) Gene ontology (GO) analyses of upregulated (D) and downregulated (E) DEGs in miR128-KD neurons (FDR < 0.05). (**F-O**) Gene Set Variation Analysis (GSVA) of the transcriptome was performed on control and miR185-KD neurons, including regulation of immune system process (F), regulation of immune response (G), cytokine production (H), chemokine production (I), learning (J), memory (K), cognition (L), positive regulation of excitatory postsynaptic potential (M), regulation of glutamatergic synaptic transmission (N), and regulation of neuronal synaptic plasticity (O). $*P < 0.05$, $**P < 0.01$, Student's t-test. DEGs: differentially expressed genes.

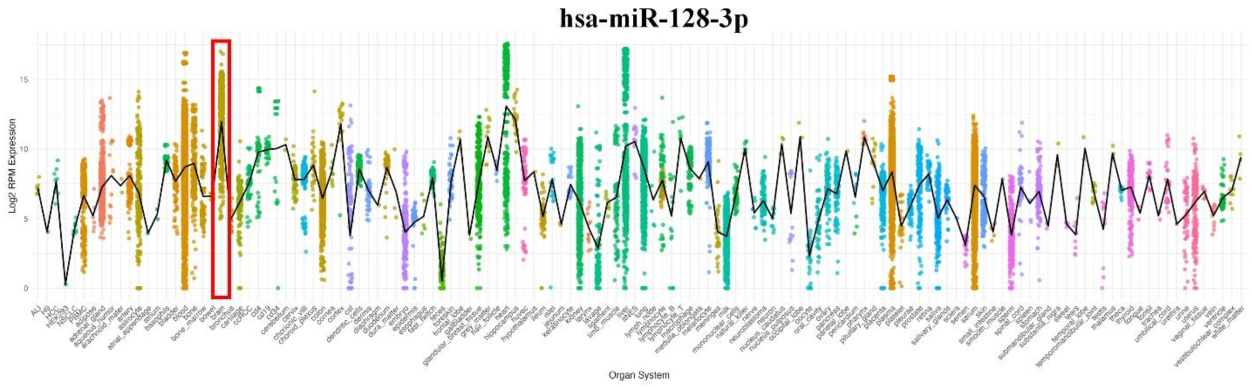


**Supplementary Figure 7. Expression profiles of hsa-miR-128-3p across different organ systems.** Data obtained from the miRNATissueAtlas2025.

### SUPPLEMENTARY TABLES

**Supplementary Table 1:** Subjects information

**Supplementary Table 2:** Differentially expressed genes in 22q11DS neurons compared with control neurons

**Supplementary Table 3:** Differentially expressed genes in 22q11DS synaptosomes compared with control synaptosomes under vehicle and TTX conditions

**Supplementary Table 4:** Differentially expressed microRNAs in 22q11DS synaptosomes compared with control synaptosomes under vehicle and TTX conditions

**Supplementary Table 5:** Differentially expressed genes in control synaptosomes following TTX treatment compared with vehicle treatment

**Supplementary Table 6:** Differentially expressed genes in 22q11DS synaptosomes following TTX treatment compared with vehicle treatment

**Supplementary Table 7:** Differentially expressed genes in hsa-miR-185-5p knockdown neurons compared with controls

**Supplementary Table 8:** Differentially expressed genes in control hsa-miR-128-3p knockdown neurons compared with controls
